# Global weight-Ranked cluster-grOWing (GROW): A Cluster-Growing Framework for Robust Functional Coordinate Network Mapping

**DOI:** 10.64898/2026.11.09.765323

**Authors:** Yu Liu, Xinglong Fu, Xin Wang, Xiaotong Wei, Hongchuan Zhang, Haoyang Dong, Yun Luo, Liyuan Lin, Qiqi Dong, Xue Zhang, Zhen Zhao, Jiaqi Liu, Shouhang Zhang, Xinjun Cao, Qianlong Ma, Mingzhen Kou, Hao Ding, Chunshui Yu, Nan Zhang, Yongqiang Zhang, Wen Qin

## Abstract

Coordinate Network Mapping (CNM) has become an important approach for translating scattered neuroimaging findings into circuit-level models of brain disorders such as Alzheimer’s disease (AD). But conventional CNM uses fixed-radius spheres and pools heterogeneous studies without accounting for confounders. We developed Global weight-Ranked cluster-grOWing (GROW), which reconstructs clusters behind reported peaks using cluster information and a reference statistical weight map, without original imaging data. We analyzed data from 521 participants and 69 published studies (82 experiments, 5,555 participants). GROW-CNM was compared with fixed-radius CNM (Fixed-CNM) in spatial fidelity, cross-study convergence, sampling stability, and sensitivity to confounders. At study level, Fixed-CNM corresponded poorly with the reference atrophy-derived functional network in AD (Spearman r = −0.24, overlap = 1.61%), and radius changes did not improve fidelity. In contrast, GROW-CNM recovered cluster morphology and generated maps closer to the reference network (r ≥ 0.85, overlap ≥ 87.30%). GROW-CNM also showed greater cross-study convergence (median peak probability, 0.99 vs. 0.67; z = 27.42, p < 0.001) and was robust to threshold/resampling (z = −169.84, p < 0.001). Fixed-CNM was associated with systematic factors; GROW-CNM was more sensitive to demographic confounders (p < 0.05, TFCE-FWE correction). Systematic confounding was attenuated by GROW-CNM; Fixed-CNM remained susceptible in core regions (p < 0.001). These findings indicate cross-study heterogeneity in coordinate-based AD network maps reflects seed definition and study-level confounders, not biology alone. Combining GROW-based seed reconstruction with confounder-adjusted population modeling may provide a robust framework for mapping disease-related brain networks and prioritizing candidate neuromodulation targets.

## Introduction

Understanding the macroscopic network architecture underlying human cognition and neuropsychiatric disorders remains a central goal of modern systems neuroscience.(Menon, 2011) A foundational advance toward this goal was Lesion Network Mapping (LNM), an approach that links local brain damage to distributed neural circuits by projecting patient-specific focal lesion locations onto normative human connectomes,(Boes et al., 2015) thereby uncovering the causal network mechanisms behind diverse neurological and psychiatric symptoms.(Fox, 2018) However, LNM typically relies on individual lesion locations, limiting its applicability when patient-specific lesion is unavailable. Evolving from Lesion Network Mapping (LNM), Coordinate Network Mapping (CNM) has rapidly become one of the most influential techniques that translates fragmented findings from neuroimaging literature—such as peak coordinates of structural atrophy,(Wang et al., 2015) functional activation,(Yarkoni et al., 2011) or localized microstructural alterations(Paquola et al., 2019)—into coherent circuit-level representations. By projecting coordinates reported across independent studies onto a normative functional connectome, CNM enables heterogeneous and often irreproducible focal findings to be reinterpreted as convergent nodes within a shared distributed network.(Taylor et al., 2021) This logic has reshaped our conceptualization of psychopathology, revealing symptom-specific circuits in disorders such as depression,(Zhukovsky et al., 2021) schizophrenia,(Wang et al., 2024) and Alzheimer’s disease,(Darby et al., 2019) and providing a principled framework for nominating neuromodulation targets.(Siddiqi et al., 2022) Alzheimer’s disease (AD) is one of the most active testbeds for CNM. Voxel-based morphometry (VBM) studies of AD report peaks of gray matter loss that are notoriously inconsistent across studies, including the medial temporal lobe (MTL), posterior cingulate/precuneus cortex (PCC), angular gyrus (AG), and lateral prefrontal cortex (LPFC) (Yang et al., 2012). Atrophy network mapping (ANM)—which leverages the normative human connectome to unify these disparate foci within a common framework—offers an appealing resolution by linking these spatially divergent atrophy peaks onto a common functional network in AD.(Tetreault et al., 2020) This network perspective is particularly valuable for identifying circuits associated with disease progression, cognitive decline, and clinical heterogeneity,(Lee et al., 2026) where pathological processes in AD extend beyond their initial site and regional atrophy patterns may reflect disruption of interconnected functional systems. Despite this promise, the reliability and specificity of CNM-derived networks relative to ANM in AD, and across different disorders remain to be systematically established.(Taylor et al., 2021) The credibility of any CNM map rests entirely on two operations that current practice treats as trivial: how a reported coordinate is converted into a seed, and how study-level maps are aggregated into a population consensus. The first issue arises within the study level. Most existing studies use reported peak coordinates as proxies for the underlying atrophy clusters and represent each coordinate as a spherical seed of arbitrary, fixed radius—typically 4-10 mm(Ji et al., 2026). This practice makes CNM computationally straightforward and facilitates the synthesis of published literature. However, this approach discards essential spatial information (i.e., size/shape) of reported clusters, treating a small focal cluster identically to a massive cortical cluster spanning multiple functional territories. Because functional connectivity varies steeply across the sub-networks,(Shen et al., 2023) imposing uniform spherical geometry introduces systematic spatial distortion. Such simplification is especially problematic in AD, in which atrophy often involves spatially extended and anatomically distributed regions.(Lam et al., 2013) Consequently, the resulting network maps may depend more on the chosen seed radius than on the spatial organization of disease-related atrophy.

A second concern arises at the population level. Consensus CNM maps are commonly generated by binarizing and overlaying study-level network maps, implicitly treating studies as exchangeable observations of a common biological circuit.(Darby et al., 2019) However, AD neuroimaging studies differ in both systematic factors (such as MRI vendor, magnetic field strength, preprocessing procedures, gray matter morphometry estimation, statistical thresholding, and so on) and sample factors (including education, sex distribution, disease severity, and cognitive status, etc.) (Abdulkadir et al., 2011; Bürkle et al., 2025; Sauty et al., 2023). These differences may influence both the location and extent of reported atrophy clusters and therefore bias the resulting network maps. Failing to account for these confounders may introduce literature-wide methodological bias into the inferred disease biology. The vulnerability of peripheral network regions to such influences is particularly important because these regions are sometimes used to interpret disease biological diversity(Lam et al., 2013).

We therefore tested three linked hypotheses. First, we posited that the fixed-radius abstraction is not a benign simplification but a source of systematic errors, such that study-level CNM maps may diverge from the true atrophy-derived network. Second, we hypothesized that recovering the original statistical cluster underlying each reported peak — using only the information contained in statistical parametric maps, without access to raw imaging data — is sufficient to restore study-level network fidelity and enhance population-level convergence. Third, we proposed that a substantial component of the between-study heterogeneity previously attributed to biological variation in AD is partially methodological, with technical and cohort-related factors exerting spatially structured effects on CNM estimates.

To test these hypotheses, we developed Global weight-Ranked cluster-grOWing (GROW), an algorithmic framework designed to reconstruct original statistical clusters from reported peak coordinates and a referential statistical parametric map topography. GROW uses globally ranked statistical evidence and competitively constrained region growing to identify cluster boundaries according to empirical statistical topography rather than a fixed-radius spherical prior. We integrated GROW into the CNM pipeline (GROW-CNM) and established a quantitative evaluation framework to compare GROW-CNM against traditional fixed-radius CNM in terms of spatial fidelity, cross-study concordance, analytical stability, and resilience to systematic and demographic confounds. An overview of the study design is provided in **Fig. 1**.

**Figure 1.**
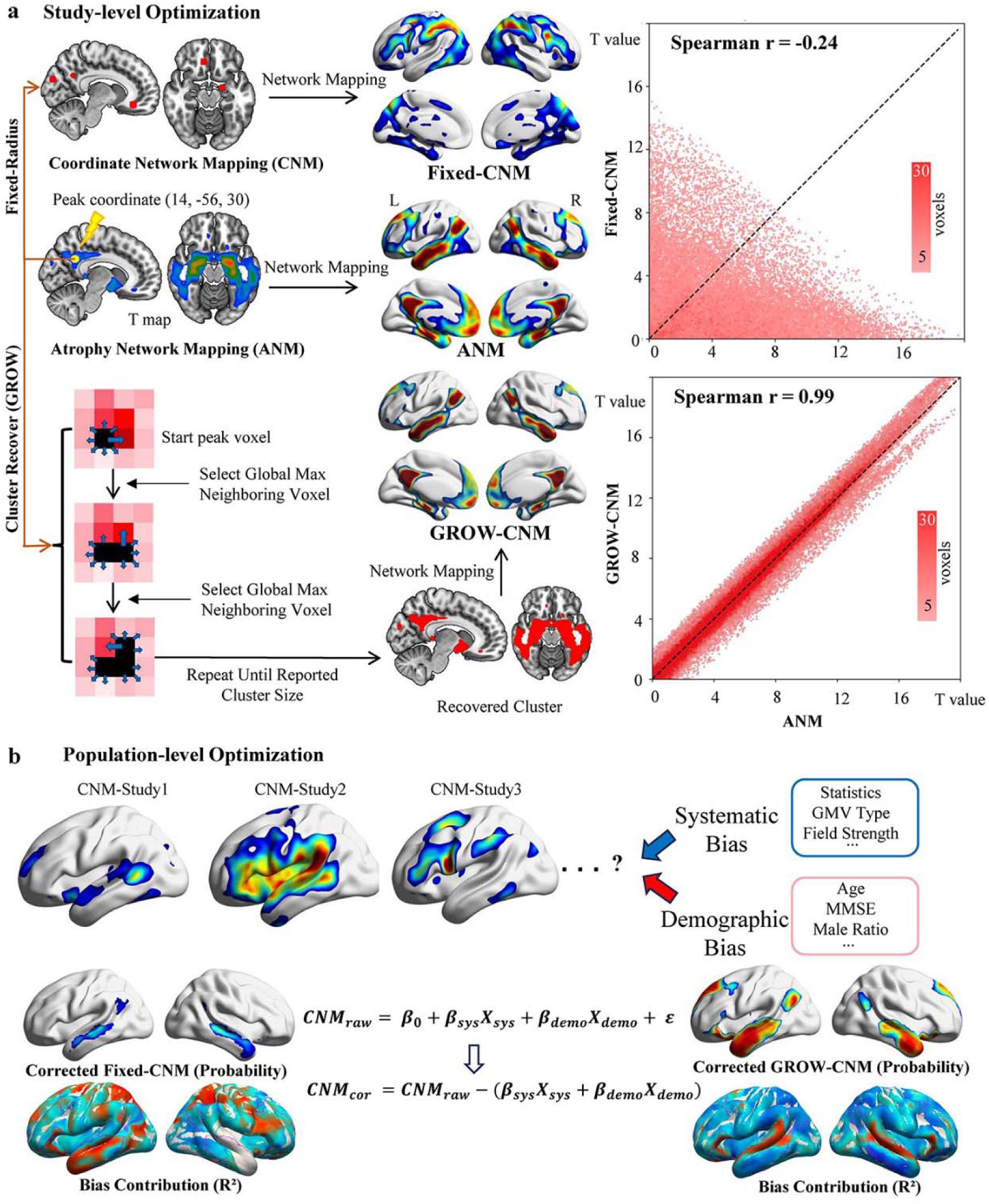
Conceptual pipeline and illustrative reconstruction of ANM, Fixed-CNM and GROW-CNM. (a) Schematic illustration of the three approaches. For ANM, the original atrophy cluster was used as the seed. For Fixed-CNM, a fixed-radius spherical seeds were generated at reported peak coordinates. For GROW-CNM, the seed was reconstructed by iterative region growing from the peak coordinate using un-thresholded statistic maps as the reference until the reported cluster size was reached. The right panels show the spatial correlation between Fixed-CNM, GROW-CNM and ANM. (b) Population-level optimization framework for confounder adjustment. Study-level CNM maps from individual experiments were integrated to generate population-level probability maps. The confounder-adjusted CNM map was obtained by removing the variance attributable to the systematic and demographic confounders. Bias contribution (R²) was quantified as the total variance contributed by systematic and demographic factors, separately. The same optimization framework was applied to both Fixed-CNM and GROW-CNM. ANM = Atrophy Network Mapping; CNM = Coordinate Network Mapping; GROW = Global weight-Ranked cluster-grOWing.

## Methods

### Participants and dataset

The local cohort comprised 42 Alzheimer’s disease (AD) patients (mean age 65.38 ± 8.47 years) and 49 normal controls (NC) (mean age 66.82 ± 6.03 years) recruited from Tianjin Medical University General Hospital between 2017 and 2021. All participants provided written informed consent, and the study protocol was approved by the Ethics Committee of Tianjin Medical University General Hospital (Approval No. IRB2020-YX-161-01). AD patients were diagnosed according to the IWG-2 criteria.(Dubois et al., 2014) Normal controls were cognitively normal (MMSE ≥ 24) and free from the same exclusion criteria. The same dataset has been used in a prior publication,(Dong et al., 2023) to which we refer readers for additional details.

ADNI datasets used in this study were obtained from the Alzheimer’s Disease Neuroimaging Initiative (ADNI) database (http://adni.loni.usc.edu). We selected AD patients and normal controls (NC) from the ADNI 1/GO/2/3 datasets with available T1-weighted structural MRI data. Diagnoses of AD were based on the criteria reported in the ADNI database, which have been described previously.(Petersen et al., 2010) We included only participants with a diagnosis of AD or NC at baseline. To minimize selection bias and control for these potential confounding variables, we performed propensity score matching (PSM) using a logistic regression model with covariates including age, sex and total intracranial volume (TIV). AD patients and NC were matched in a 1:1 nearest-neighbor algorithm without replacement. After matching, 215 AD patients and 215 NC were retained for subsequent analyses. Detailed information was provided in **Supplementary Table 1**.

The local TMUGH cohort (42 AD, 49 NC) and the matched ADNI cohort (215 AD, 215 NC) were combined into a merged dataset, yielding a total of 257 AD patients and 264 NC. This combined dataset was used for GMV statistical parametric mapping for ANM and CNM estimation. Detailed information was provided in **Supplementary Table 2**.

T1-weighted 3D structural MRI (sMRI) data for the TMUGH cohort were acquired on a 3.0-Tesla Discovery MR750 scanner (General Electric, Milwaukee, WI) using a brain volume (BRAVO) sequence. Detailed acquisition parameters have been described previously.(Dong et al., 2023) The sMRI data of ADNI cohort were acquired using 3D T1-weighted magnetization prepared rapid acquisition gradient-echo (MPRAGE) imaging sequences. The detailed acquisition information was available at the ADNI website (http://adni.loni.usc.edu/methods/mri-tool/mri-analysis/).

### MRI data preprocessing

All T1-weighted structural MRI data were preprocessed using the CAT12 toolbox (http://www.neuro.uni-jena.de/cat), an extension of SPM12 (https://www.fil.ion.ucl.ac.uk/spm/software/spm12) running on MATLAB. The preprocessing pipeline included: (1) bias correction to correct B1 field inhomogeneity-induced signal variation across voxels; (2) tissue segmentation into gray matter (GM), white matter, and cerebrospinal fluid; (3) spatial normalization of the GM tissue into Montreal Neurological Institute (MNI) space using the DARTEL algorithm, with modulation by Jacobian determinants to preserve absolute GM volume (GMV), resampled to 1.5-mm isotropic voxels; and (4) spatial smoothing with an 8-mm full-width at half-maximum (FWHM) Gaussian kernel.

### Literature search and study selection

In addition to the individualized GMV analysis, a systematic literature search was also conducted on November 6, 2024, using the Web of Science and PubMed databases. The search syntax was: ((Alzheimer’s Disease) OR (Alzheimer Disease)) AND ((voxel-based morphometry) OR (VBM) OR (GMV) OR (Gray Matter Volume) OR (Grey matter volume) OR (GMD) OR (Gray Matter Density) OR (Grey matter density) OR (GMC) OR (gray matter concentration) OR (grey matter concentration)), restricted to title/abstract fields.

The study selection process followed the Preferred Reporting Items for Systematic Reviews and Meta-Analyses (PRISMA) guidelines (**Figure 2**).(Page et al., 2021) specifically, studies were included if they met the following criteria: (1) reported comparisons between AD patients and normal controls; (2) reported results with both peak coordinates (x, y, z) and cluster size information; and (3) were peer-reviewed and published in English; (4) When overlapping data were identified across studies, the one with the largest sample size was retained. A total of 3,905 records were identified from Web of Science and PubMed. After duplicate removal and screening, 69 studies (82 independent experiments) met the inclusion criteria for subsequent CNM analyses.

**Figure 2.**
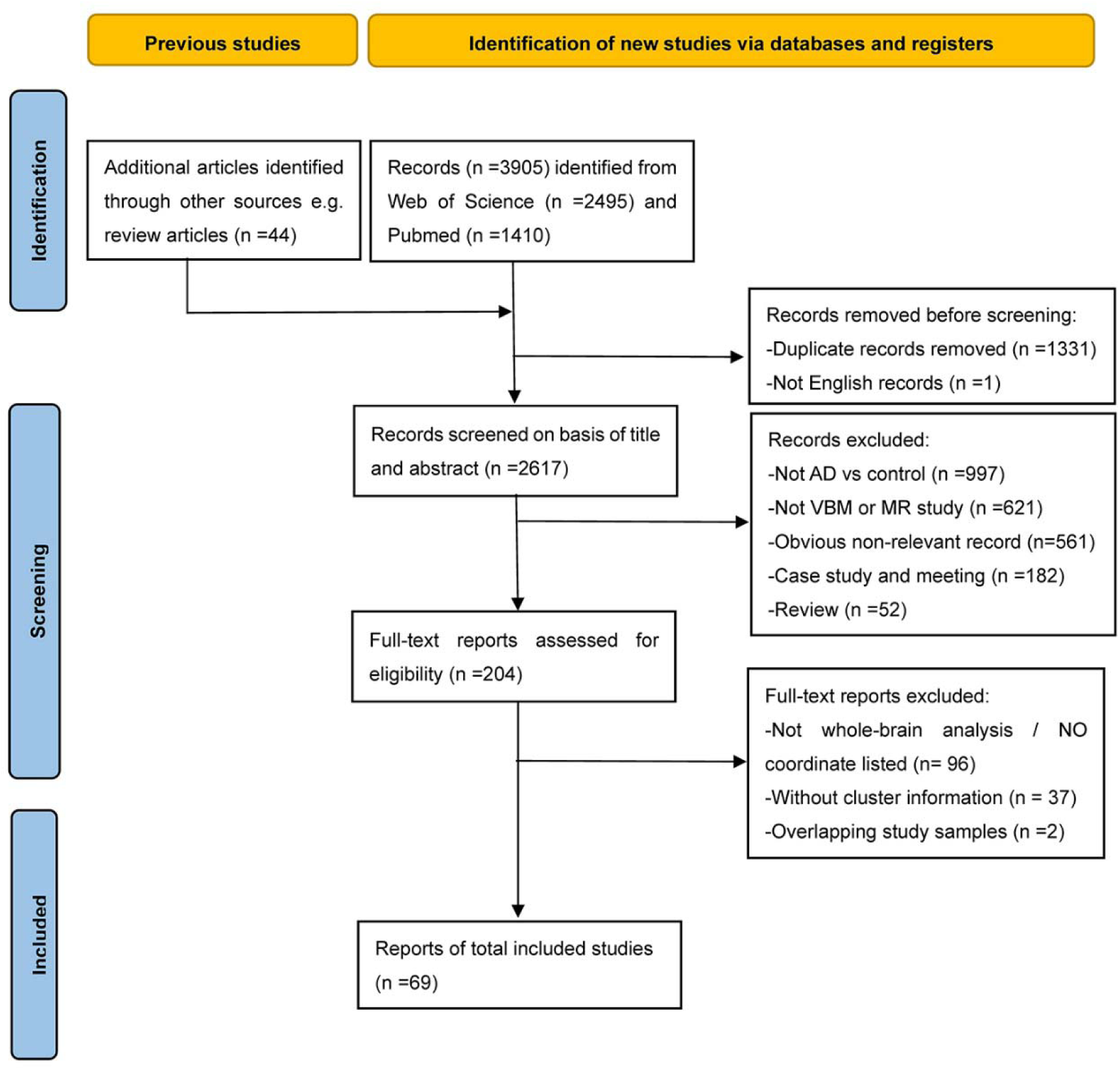
PRISMA flow diagram of the systematic literature search and study selection. Flowchart summarizing the identification, screening, eligibility assessment, and inclusion of studies.

### Definition of potential literature-level confounders

We extracted a set of potential literature-level confounding variables from each included study, covering both systematic and sample (demographic/clinical) characteristics factors.

Systematic factors included: MRI vendor (categorized as Siemens, GE, Philips, vs. others), field strength (categorized as < 3T vs. ≥ 3T), GMV type (absolute GMV vs. GM density/concentration), smoothing kernel (FWHM), statistical model (parametric vs. non-parametric tests), and correction strategy for multiple comparisons (voxel-level, cluster-level, TFCE-FWE, or uncorrected). Sampling factors included: mean age, sex (male ratio), mean education years, and mean MMSE score.

For variables with missing data, continuous variables were imputed using the median value across available studies, while categorical variables were imputed using the mode. Detailed coding rules for each variable, including category consolidation, are provided in **Supplementary Table 3**, and the complete variables information after imputing is provided in **Supplementary Table 4**.

### Creation of local atrophy statistical maps and coordinate extraction

For the MRI dataset, voxel-wise two-sample t-tests were performed to compare GMV between AD patients and normal controls using SPM12. For the TMUGH and ADNI cohorts, age, sex, education years, and total intracranial volume (TIV) were included as nuisance covariates. For the merge cohort, an additional covariate of dataset source (TMUGH vs. ADNI) was included to account for inter-site heterogeneity.

Two types of statistical maps were generated: (1) uncorrected t-statistic maps, which were retained as reference maps for subsequent seed reconstruction using GROW algorithms; and (2) thresholded maps at voxel-level FWE correction (p < 0.05), which were used as inputs for atrophy lesion network mapping (ANM) and cluster coordinates reporting. For each significant cluster of the thresholded maps, we recorded the peak Montreal Neurological Institute (MNI) coordinates and the cluster size (in mm^3^), which were subsequently used as inputs for the Fixed-CNM and GROW-CNM procedures.

### Atrophy network mapping

For each study, the ANM was performed using a standardized pipeline.(Fox, 2018) The thresholded atrophy maps (voxel-level FWE-corrected p < 0.05) derived from the GMV analyses were used as seed regions. The normative functional connectome was constructed using preprocessed functional MRI data from 100 healthy subjects at the TMUGH site, a subset of Chinese Imaging Genetics (CHIMGEN) cohort.(Xu et al., 2020) The standard preprocessing pipeline included: distortion correction, slice timing, realign, DARTEL normalization, nuisance regression (24 Friston motion parameters, spikes with Power FD>0.5, global, white matter and cerebrospinal fluid signals.), and filtering (0.01-0.08 HZ). Then, the functional connectivity (FC) map of each seed per participant to every other brain voxel was calculated using the Pearson’s correlation coefficients between the mean time series of the seed and ones of each other voxels and was Fisher’s z-transformed. The resulting individual FC maps were smoothed with a 6-mm FWHM Gaussian kernel. Finally, a one-sample t-test across normative participants was then performed to generate study-level ANM t-statistic maps for each dataset (**Supplementary Fig. 1**).

### Fixed-CNM construction

For the fixed-radius CNM (Fixed-CNM) approach, spherical seeds were generated centered at each reported peak coordinate from the included VBM studies. A 6-mm radius was selected as the primary sphere size, which has been commonly adopted in CNM studies to balance spatial specificity and functional connectivity sensitivity(Taylor et al., 2021; Zhukovsky et al., 2021). To assess the robustness of our findings, we additionally generated sphere per coordinate with 4-mm and 8-mm radii as sensitivity analyses. All coordinate spheres within one study were then merged into one single combined binary seed mask. Finally. for each study, the mean resting-state time series of the combined seed were extracted from the normative CHIMGEN connectome, and the study-level fixed-CNM map was calculated following the same procedure as described in the ANM section.

### GROW algorithm and GROW-CNM construction

To reconstruct the spatial extent of the cluster associated with each reported coordinate, we developed the Global weight-Ranked Cluster-groWing (GROW) algorithm (Fig. 1a). GROW adapts the seeded region-growing framework(Adams and Bischof, 1994), but replaces its local neighbor search with a global best-first strategy(Falcao et al., 2004). The algorithm takes three inputs: (1) the reported peak coordinate in MNI space, (2) the reported cluster size, and (3) a reference weight map, here the unthresholded t-statistic map from the AD case–control comparison. Growth starts at the voxel containing the peak coordinate. GROW keeps a queue of every voxel adjacent to the current cluster. At each iteration, it adds the queued voxel with the highest weight (t-value), wherever that voxel lies on the cluster boundary. Growth stops when the cluster reaches the reported size. The output is a binary mask that matches the reported cluster size and follows the strongest disease-related effects in the weight map, such as atrophy in AD.

Then all clusters of one study were then merged into one single binary seed mask. Finally, the GROW-based CNM (GROW-CNM) for each study were derived using the same normative CHIMGEN connectome and the procedure as described in the ANM section.

### Study-level performance comparison

To evaluate study-level spatial fidelity, we compared Fixed-CNM and GROW-CNM with the reference ANM in both the TMUGH and merged datasets. First, we computed the Spearman rank correlation between each CNM-derived unthresholded connectivity map (t-statistic maps) and the corresponding ANM map. Second, we quantified the spatial overlap between each CNM-generated thresholded connectivity mask (A) and the reference ANM connectivity mask (B) (t threshold > 5) using the Dice coefficient (Formula [1]):

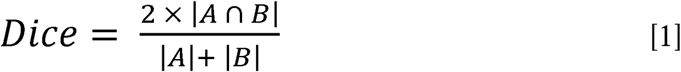

### Population-level performance comparison

We generated population-level probability maps for Fixed-CNM and GROW-CNM by binarizing each study-level functional connectivity map at a prespecified study-level t-threshold and calculating the proportion of studies with suprathreshold connectivity at each voxel. To assess spatial agreement between methods, we thresholded both probability maps at 0%, 25%, 50%, and 75% and calculated the proportion of suprathreshold voxels in the GROW-CNM map that overlapped with the Fixed-CNM map. We also computed the voxel-wise Spearman correlation between the unthresholded probability maps and visualized their relationship using a scatter plot.

Besides, to assess consistency across study-level thresholding criteria, we generated probability maps using t-thresholds of 3.1, 5, and 7. We then compared each pair of maps using the intraclass correlation coefficient (ICC) and the Dice coefficient, with a 50% probability threshold applied for the latter.

To assess robustness across studies, we generated 1,000 bootstrap samples by sampling studies with replacement, with each sample matching the original number of studies. For each sample, we recomputed the probability maps and recorded the peak probability across the brain. We plotted the distribution of peak probabilities and calculated the mean probability and coefficient of variation (CV) at each voxel across bootstrap samples. For GROW-CNM, we fitted a power-law model to the voxel-wise relationship between mean probability and CV using nonlinear least squares and evaluated goodness of fit using R². Finally, we compared peak probabilities and CVs in core regions (probability>50%) between Fixed-CNM and GROW-CNM using Wilcoxon signed-rank test (P<0.05, Bonferroni correction).

### The effect of single confounders on population-level CNM

We assessed the effects of individual systematic and demographic factors using the voxel-wise general linear model (GLM), fitted separately for Fixed-CNM and GROW-CNM. At each voxel, study-level CNM values (t values) were modeled as a function of one factor at a time. Factors with more than two categories, such as MRI vendor, were modeled as multilevel categorical variables. Multiple comparisons across voxels were controlled using threshold-free cluster enhancement (TFCE) with family-wise error correction (p < 0.05; 5,000 permutations). Uncorrected results (p < 0.05) were also displayed for exploratory purposes.

### Multi-confounders adjustment for population-level CNM and performance comparison

To account for potential confounding effects identified in the single-factor GLM analyses, we used multivariable regression to generate confounder-adjusted population-level CNM probability maps for Fixed-CNM and GROW-CNM. All factors showing spatially clustered effects at an uncorrected (p < 0.05) in the single-factor analyses were included in the adjustment models.

We used a hierarchical adjustment strategy: the first model included systematic factors only, whereas the second included both systematic and demographic factors. This produced three sets of population-level probability maps for each method: unadjusted, systematic-adjusted, and fully adjusted.

For each method, we calculated voxel-wise R² values for the systematic-only and full models to quantify the proportion of variance explained by the included confounders (formula [2]):

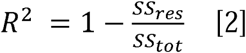

where *SS_res_* and *SS_tot_* denote the residual and total sums of squares, respectively. Specifically, we calculated R² for the systematic-only model and the full model (systematic + demographic), separately for Fixed-CNM and GROW-CNM. At each voxel, the full model was specified as formula [3]:

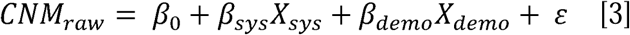

where *X_sys_* and *X_demo_* represent the systematic and demographic confounders, respectively, and β*_sys_* and β_demo_ are their corresponding coefficients estimated from the multivariable GLM. The intercept β_0_ represents the expected CNM value when continuous confounders equal zero and categorical confounders are at their reference levels.

The confounder-adjusted CNM values for each study were obtained by subtracting the estimated contributions of the included confounders (formula [4]):

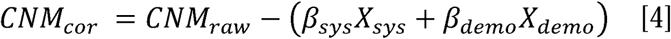

For the systematic-only model, the demographic term was omitted. Adjustment was performed separately for Fixed-CNM and GROW-CNM.

To assess how adjustment affected the spatial extent of each map, we binarized the adjusted and unadjusted probability maps at a threshold of 0.5. We then calculated the percentage of suprathreshold voxels in the unadjusted map that remained suprathreshold after adjustment. This measure was computed for both adjustment models and both CNM methods.

To compare the distributions of voxel-wise R² values, we compared R² distributions across the following dimensions using non-parametric two-sided tests: (1) Between-method comparisons: GROW-CNM versus Fixed-CNM within the core region, separately for the systematic-only and full models, using the Wilcoxon signed-rank test. (2) Within-method comparisons: core versus non-core regions, separately for each method and model, using the Mann–Whitney U test. The core region was defined as voxels with probability ≥ 0.5 in the unadjusted CNM maps (P < 0.05, Bonferroni correction).

### Sensitivity analyses

To evaluate the robustness of our findings, we performed three sensitivity analyses: (1) Seed radius: For Fixed-CNM, we varied the seed radius (4, 6, and 8 mm) to assess sensitivity to seed size. (2) Weight map: For GROW-CNM, we used three GMV statistical maps as weights—the TMUGH-derived, ADNI-derived, and merged t-maps—and compared the resulting connectivity and population-level probability maps. (3) Study-level threshold: For both methods, we repeated the pipeline using multiple voxel-level t-thresholds (e.g., 3.1, 5.0, and 7.0) to determine whether between-method differences depended on the threshold.

## Results

### Participant and literature characteristics

The demographic and clinical characteristics of the ADNI and merged samples are summarized in **Supplementary Tables 1 and 2**, respectively. Age, sex distribution, and years of education did not differ significantly between the AD and normal control (NC) groups in either sample (all p > 0.05). As expected, patients with AD had lower MMSE scores than NCs in both samples (both p < 0.001).

A total of 69 studies comprising 82 independent experiments met the inclusion criteria for CNM analyses (**Fig. 2**). These experiments included 5,555 participants: 2,685 patients with AD (mean age, 71.90 years; 1,228 males and 1,457 females) and 2,870 NCs (mean age, 69.77 years; 1,329 males and 1,541 females). The extracted study-level systematic and demographic variables are summarized in **Supplementary Table 4**. Missing data proportions were 4.88% for sex ratio, 30.49% for years of education, 21.95% for MMSE, 9.76% for MRI vendor, 1.22% for smoothing kernel, and 1.22% for statistical model.

### The performance of GROW method in recovering cluster morphology

The spatial distribution of GMV atrophy in AD is shown in **Supplementary Fig. 2**, and the cluster results for the TMUGH and merged datasets are presented in **Supplementary Tables 3 and 4**, respectively.

For the TMUGH dataset, GROW-reconstructed clusters closely matched the original atrophy clusters when using the TMUGH-derived t-map as the weight map (Dice = 0.95). When the ADNI or merged t-map was used as the reference, GROW recovered nearly half of the TMUGH atrophy clusters (0.48 and 0.58, respectively) (**Supplementary Fig. 3**). For the merged dataset, GROW recovered the atrophy cluster morphology even better (Dice= 0.97, 0.94, and 0.72 using the merged-, ADNI-, and TMUGH-derived weight maps, respectively) (**Supplementary Fig. 4**).

### GROW-CNM improves study-level correspondence with ANM

The study-level spatial distribution of each CNM connectivity map and the ANM is presented in **Fig. 3a (**thresholded t>5). Study-level CNM maps were compared with the corresponding ANM reference using voxel-wise Spearman correlation (unthresholded t-map, **Fig. 3b**) and spatial overlap (t>5, **Fig. 3c**).

**Figure 3.**
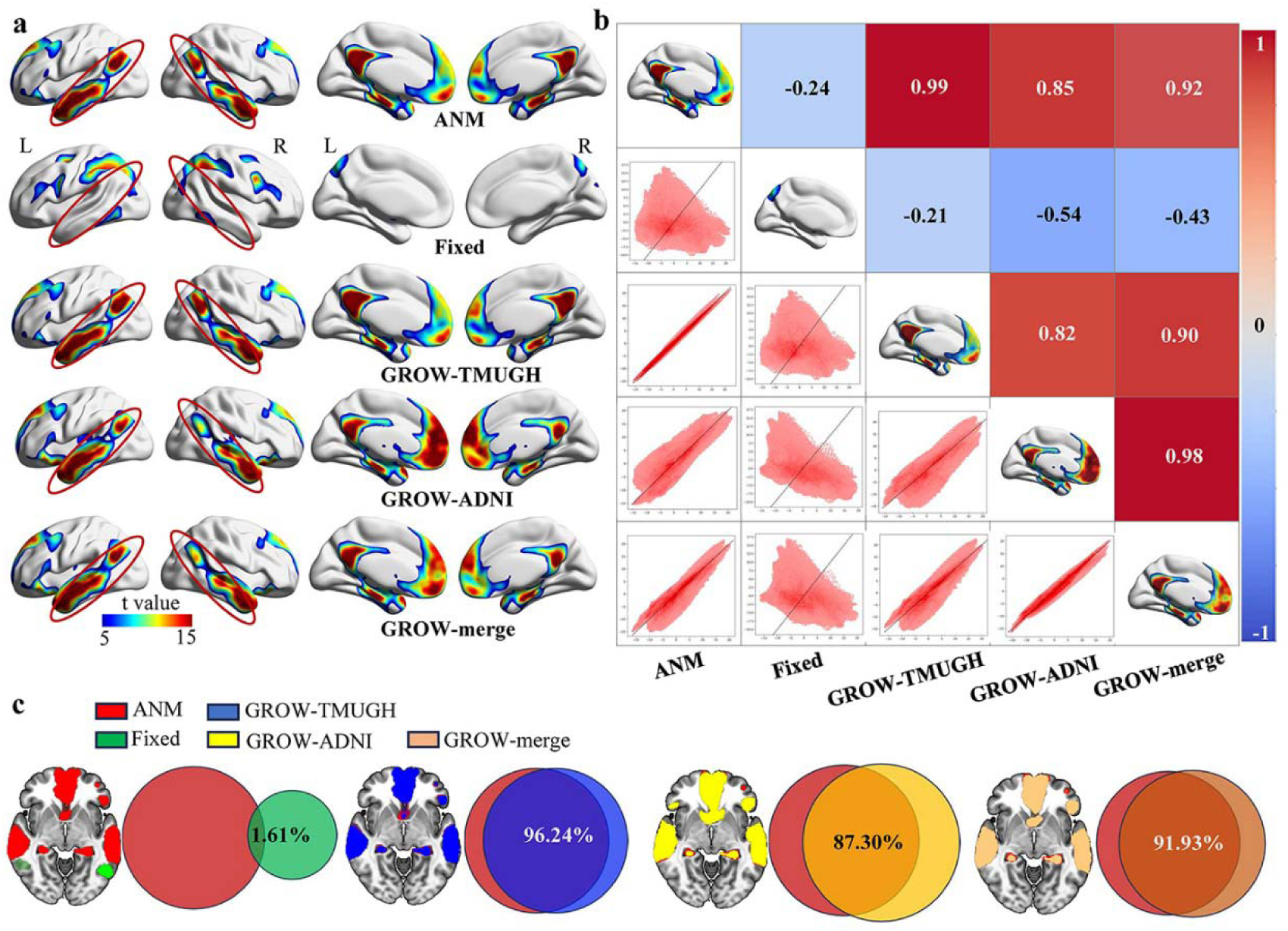
Study-level comparison of ANM, Fixed-CNM and GROW-CNM. **(a)** study-level thresholded ANM, Fixed-CNM and GROW-CNM reconstructed using reference weight map from TMUGH (GROW-TMUGH), ADNI (GROW-ADNI), and merged dataset (GROW-merge), respectively (t >5). **(b)** Spearman correlation between each pair of network maps. Numbers indicate Spearman correlation coefficient. **(c)** Overlap between ANM and different types of CNM maps (t >5). LNM = Lesion Network Mapping; GROW = Global weight-Ranked cluster-grOWing; ADNI = Alzheimer’s Disease Neuroimaging Initiative; TMUGH = Tianjin Medical University General Hospital.

In the TMUGH dataset, fixed-CNM showed poor correspondence with ANM (r = −0.24; overlap=1.61%). Fixed-CNM maps generated using 4-, 6-, and 8-mm seeds were highly correlated (r >0.95), excluding the possibility of seed radius on the poor correspondence with ANM (**Supplementary Fig. 5**). In contrast, GROW-CNM showed greater correspondence across all three reference weight maps: TMUGH-derived (r = 0.99; overlap = 96.24%), merged-derived (r=0.92, overlap=91.93%), and ADNI-derived (r=0.85, overlap=87.30%). A similar pattern was observed in the merged dataset: Fixed-CNM showed poor correlation with ANM (r = −0.10), whereas GROW-CNM showed higher correlations using the merged-, ADNI-, and TMUGH-derived reference weight maps (r = 0.98, 0.99, and 0.42, respectively; **Supplementary Fig. 6**).

### GROW-CNM shows greater cross-study spatial concordance and stability than Fixed-CNM

We compared population-level probability maps generated by Fixed-CNM and GROW-CNM. Fixed-CNM produced a diffuse map with generally low probabilities, whereas GROW-CNM showed higher probabilities in spatially focused network regions (**Fig. 4a**). Across voxels, the median probability was 0.293 (IQR = 0.634) for GROW-CNM and 0.110 (IQR = 0.183) for Fixed-CNM. GROW-CNM also retained more voxels at higher probability thresholds (**Fig. 4a, Supplementary Fig. 7**). Moreover, the proportion of suprathreshold overlapped voxels between Fixed-CNM and GROW-CNM maps decreased from 99.3% (unthresholded) to 66.5% (probability>25%), 11.3% (probability>0.50) and disappeared at probability of 75% (**Fig. 4a**). Although the two unthresholded probability maps were strongly correlated (r = 0.86; **Fig. 4b**), most voxels had higher probabilities in GROW-CNM than in Fixed-CNM.

**Figure 4.**
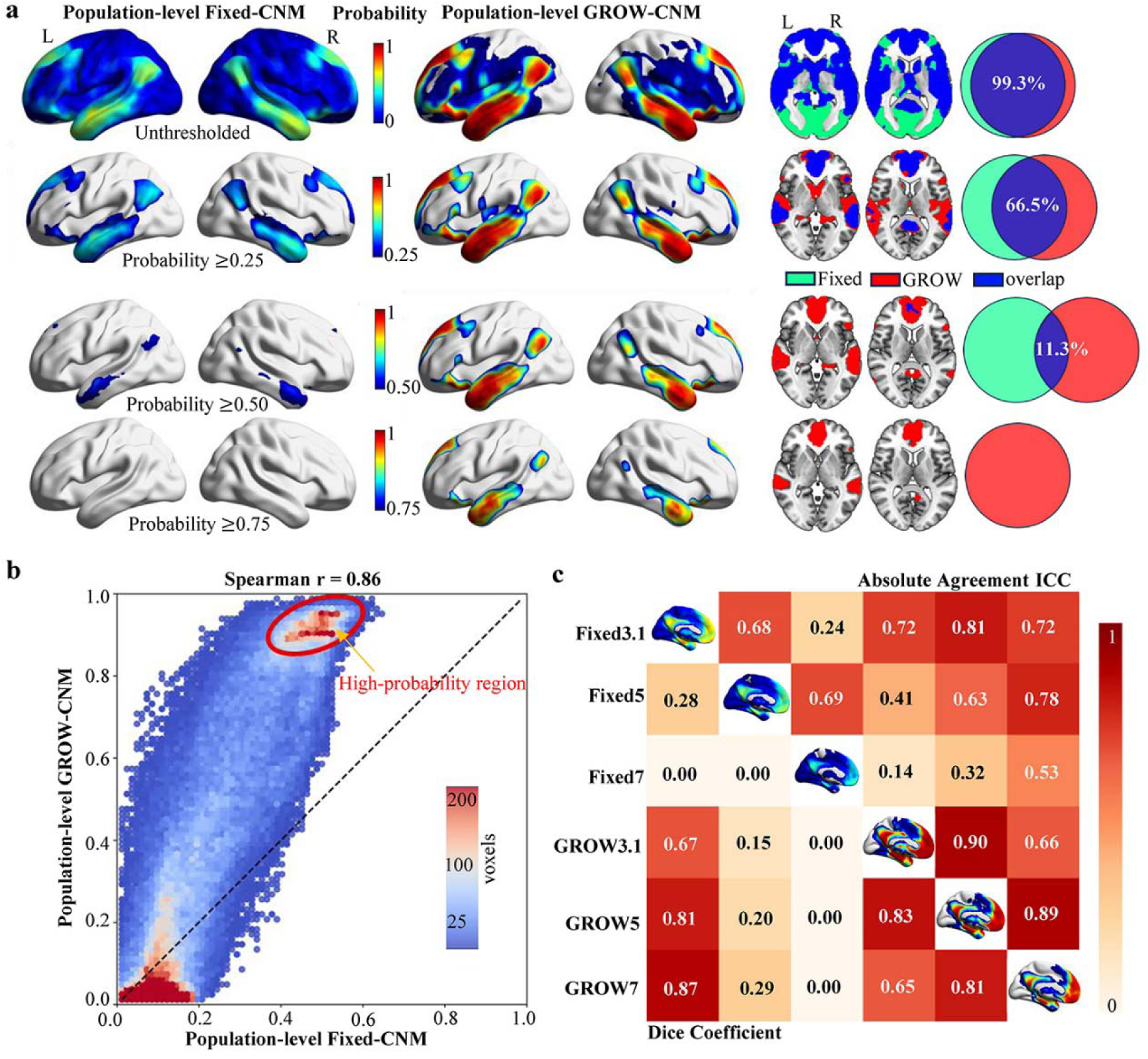
Population-level probability maps of Fixed-CNM and GROW-CNM without confounder adjustment. **(a)** Probability maps for Fixed-CNM and GROW-CNM at thresholds of 0%, 25%, 50%, and 75% (left) and their spatial overlap at each threshold (right). (b) Voxel-wise scatter plot of probability values between Fixed-CNM and GROW-CNM (Spearman’s r = 0.86). (c) Pairwise spatial consistency of CNM maps generated using study-level thresholds of t = 3.1, 5, and 7, assessed by Dice coefficients (lower triangle) and intraclass correlation coefficients (ICCs; upper triangle). CNM, Coordinate Network Mapping; GROW, Global weight-Ranked cluster-grOWing.

Across study-level thresholds of t = 3.1, 5, and 7, GROW-CNM showed greater within-method consistency than Fixed-CNM (mean ICC = 0.82 vs. 0.54; **Fig. 4c** and **Supplementary Fig. 8**), indicating that GROW-CNM was less sensitive to study-level threshold.

Bootstrap analysis with 1,000 resamples was used to assess CNM sampling stability (**Fig. 5**). Compared with Fixed-CNM, GROW-CNM showed higher mean probabilities (**Fig. 5a**) and lower coefficients of variation (CVs) in core network regions than Fixed-CNM (**Fig. 5b-c**). Across bootstrap iterations, peak probabilities were higher for GROW-CNM (median = 0.988, IQR = 0.012) than for Fixed-CNM (median = 0.671, IQR = 0.061; z = −27.42, p < 0.001; **Fig. 5d**). The lower CVs and narrower IQR of high peak probabilities are consistent with greater sampling stability of GROW-CNM. Within GROW-CNM, mean probability and CV followed a strong power-law negative relationship (R^2^ = 0.99; **Fig. 5e**), with higher-probability regions showing lower relative variability.

**Figure 5.**
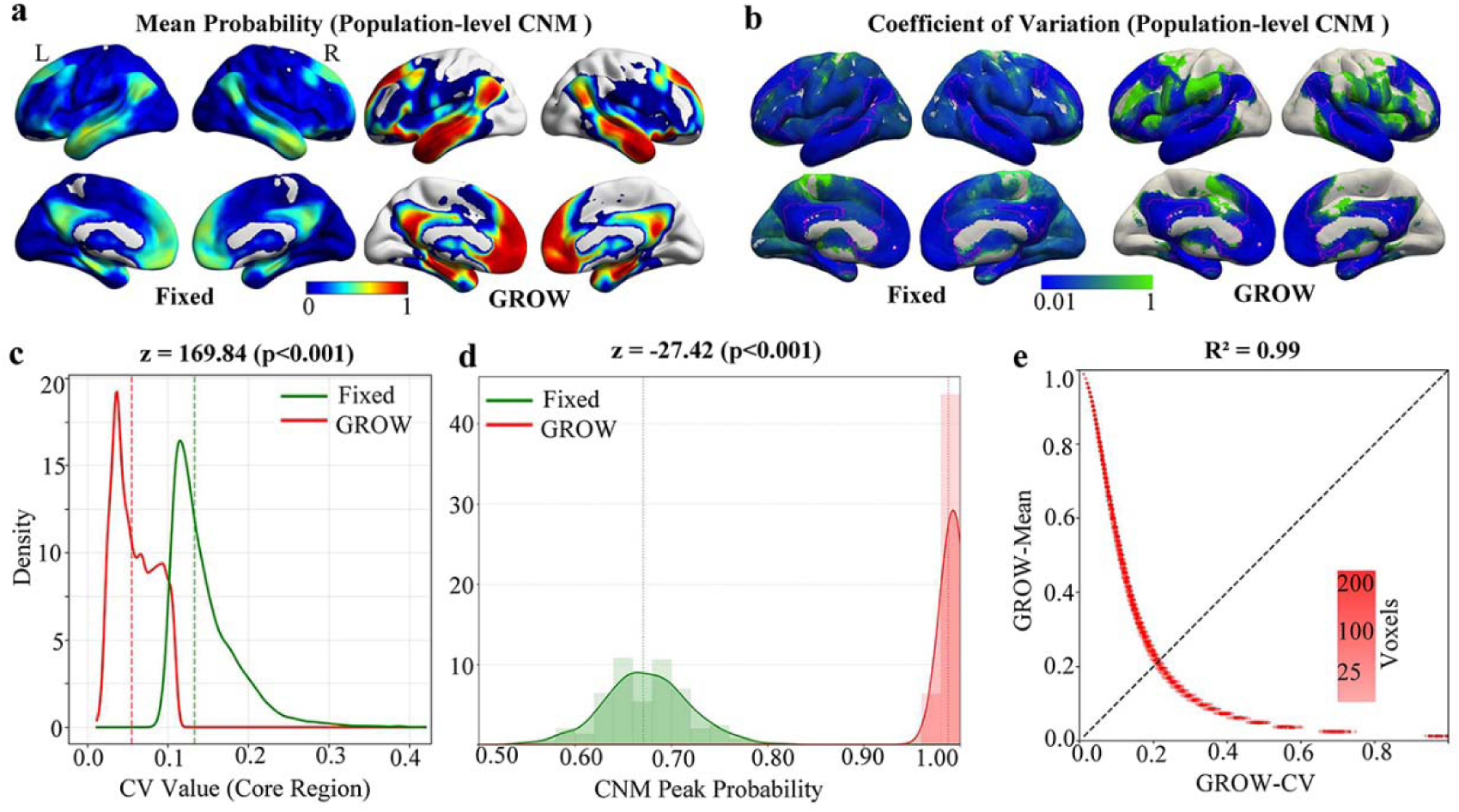
Bootstrap assessment of sampling stability for Fixed-CNM and GROW-CNM. (a) Mean probability maps across 1,000 bootstrap iterations. (b) Coefficient of variation (CV) maps across iterations; purple outlines indicate core regions. (c) Comparison of CVs within core regions between methods. (d) Comparison of peak probabilities across iterations between methods. (e) Relationship between mean probability and CV for GROW-CNM. Comparisons were performed using the Wilcoxon signed-rank test. CNM, Coordinate Network Mapping; GROW, Global weight-Ranked cluster-grOWing.

**Figure 6.**
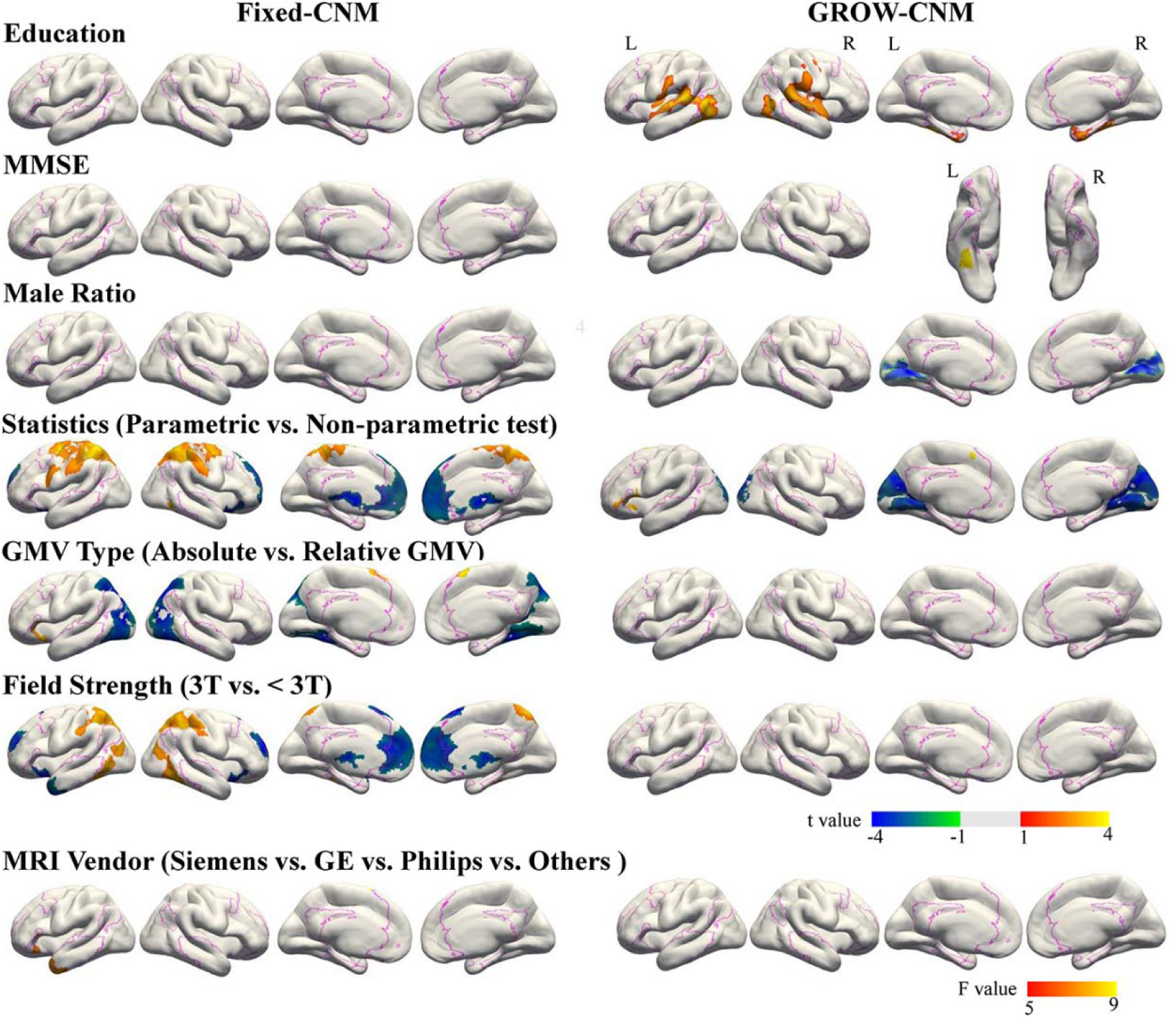
Effects of individual systematic and demographic factors on Fixed-CNM and GROW-CNM. Nonparametric voxel-wise general linear models assessed associations between each factor and study-level GROW-CNM maps. T- or F-statistic maps are shown according to factor type; only significant results are displayed (p < 0.05, TFCE-FWE corrected). In T-statistic maps, warm and cool colors indicate positive and negative associations, respectively. Purple contours delineate the core network (GROW-CNM probability > 50%). CNM = Coordinate Network Mapping; GROW = Global weight-Ranked cluster-grOWing; MMSE = Mini-Mental State Examination.

### Effects of systematic and demographic factors on Population-Level CNM

For Fixed-CNM, voxel-wise GLMs identified significant associations with four methodological factors (**Supplementary Table 7**). The statistical model (parametric vs. non-parametric) showed positive effects in the posterior parietal gyri (PPG) and somatomotor cortex, as well as negative effects in the medial prefrontal cortex (MPFC) and thalamus. GMV type (relative vs absolute GMV) showed widespread effects, including positive associations in the bilateral superior frontal gyri and negative associations in the PPG and visual association cortex. Field strengths (3.0T vs. <3T) was positively associated with the bilateral posterior parietal gyri (PPG) and occipital temporal juncions (OTJ), but negatively associated with the MPFC, thalamus, and temporal pole (TP). MRI vendors (Seimens. GE. Philips. vs others) showed significant positive effects in the left inferior frontal gyrus (IFG) and TP. There showed no statistical significant association between demographic factors (age, male ratio, education,and MMSE) and Fixed-CNM (P<0.05, TFCE-FWE correction).

For GROW-CNM, one methodological factor (statistical model) and three demographic factors (education, MMSE score, and male ratio) showed significant associations (**Supplementary Table 8**). The statistical model was negatively associated the GROW-CNM of medial occipital cortex (MOC) and positively associated with the IFG. Education exerted positive effects in the superior temporal gyrus (STG), OTJ, secondary somatosensory cortex, posterior insula, and medial temporal cortex (MTC). MMSE score showed positively effects in the fusiform gyrus, whereas male ratio manifested negative associations in the GROW-CNM of MOC (P < 0.05, TFCE-FWE correction).

When a more lenient threshold was used (p<0.05, uncorrected), all candidate systematic and demographic factors were associated with both Fixed-CNM and GROW-CNM (**Supplementary Figures 9-10**). Overall, compared with Fixed-CNM, GROW-CNM was less sensitive to systematic factors and more sensitive to demographic factors.

### Performance of multi-confounders adjustment for population-level CNM

Population-level Fixed-CNM and GROW-CNM 50% probability maps adjusted for systematic confounders and for both systematic and demographic confounders are shown in Figure 7a and Figure 7b. For Fixed-CNM, the map adjusted for systematic factors retained only 55.6% overlap with the unadjusted map, with the overlap further decreasing to 50.2% after full adjustment. In contrast, GROW-CNM retained 93.1% and 92.3% overlap after systematic and full adjustments, respectively, demonstrating that its spatial topography was substantially more resilient to confound removal than that of Fixed-CNM. The spatial distributions of proportion of variance (R^2^) explained by confounders are shown in Figure 7c. Compared with Fixed-CNM, GROW-CNM exhibited a significantly lower proportion of variance explained by either systematic or combined confounders, both across the whole brain and within core CNM regions (probability > 50%), demonstrating greater overall robustness against confounders in Fig 7d. Furthermore, GROW-CNM showed lower confounder-explained variance in core regions than in non-core regions, whereas this pattern was reversed in Fixed-CNM (Figure 7e and Supplementary Figure 11). This indicates that confounding effects in GROW-CNM were predominantly confined to peripheral, low-probability regions, whereas Fixed-CNM suffered disproportionately greater confounding in core regions.

**Figure 7.**
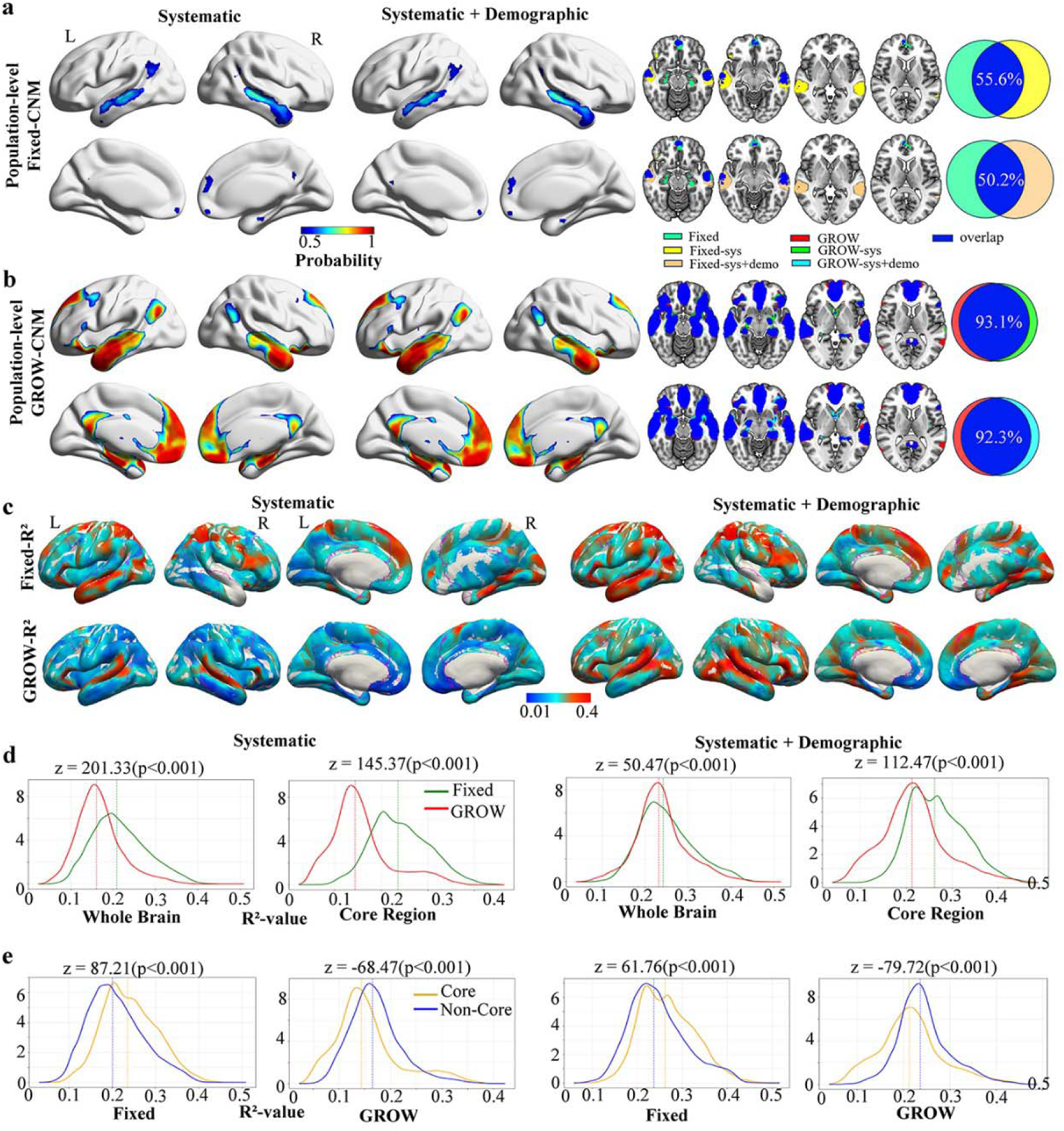
Multi-confounders adjustment for population-level Fixed- and GROW-CNM. Population-level Fixed-CNM(a) and GROW-CNM(b) 50% probability maps adjusted for systematic confounders (left panel) and both systematic and demographic confounders (middle panel). The right panel shows their overlap with unadjusted maps. (c) Variance (R²) explained by systematic **confounders** (left panel) and s both systematic and demographic confounders (right panel). Purple contours delineate the core network (GROW-CNM probability > 50%). (d) Comparison of confounder-explained variance between Fixed-CNM and GROW-CNM across the whole brain and within core regions. (e) Comparison of confounder-explained variance between core and non-core regions within each method. CNM = Coordinate Network Mapping; GROW = Global weight-Ranked cluster-grOWing.

## Discussion

This study evaluated two foundational yet largely unexamined methodological assumptions in CNM: the simplification of coordinate seeds into fixed-radius spheres(Ji et al., 2026) and the pooling of heterogeneous studies without adjustment for potential confounders(Abdulkadir et al., 2011). Using atrophy network mapping (ANM) derived directly from AD imaging data as a reference, we found that fixed-radius CNM showed poor study-level correspondence with ANM and could even reverse the direction of study-level network profile. In contrast, CNM based on the reconstruction of native cluster morphology through global weight-ranked cluster-growing (GROW) yielded higher correspondence with ANM, greater population-level cross-study convergence, and improved resampling stability. Moreover, a substantial proportion of the between-study variability in CNM also appeared to be influenced by both systematic and demographic confounders. These biases were substantially attenuated by GROW-CNM, whereas Fixed-CNM remained confounder-susceptible even in core regions. Together, these findings indicate that seed geometry and study-level systematic and demographic heterogeneity can influence CNM-derived networks and suggest that GROW-based cluster reconstruction and confounders adjustment may improve the robustness of circuit synthesis.

We found that fixed-radius CNM showed poor correspondence with image-derived ANM in both AD datasets, which points to a conceptual problem rather than a parameter-tuning problem. A coordinate is a spatial summary of a statistical finding rather than a complete representation of its underlying cluster(Eickhoff et al., 2012). The cluster behind a reported peak varies in extent, volume and shape depending on disease severity, statistical threshold and anatomical location(Taylor et al., 2021; Yang et al., 2012). A uniform fixed-radius sphere (e.g., 4–10 mm) may truncates large,(Ji et al., 2026) sprawling association-cortex clusters while overinflating tiny focal peaks. Given that functional connectivity varies across cortical and network boundaries(Schaefer et al., 2018; Shen et al., 2023; Yeo et al., 2011), these geometric mismatches can corrupt the connectivity profile. Our data support this explanation. Fixed-CNM maps generated with 4-, 6- and 8-mm seeds were almost identical (r > 0.95), yet all corresponded poorly with ANM (r = −0.24 in TMUGH; r = −0.10 in the merged dataset). The bias therefore comes from collapsing clusters into points, not from choosing the wrong radius, and thus cannot be readily resolved by changing the sphere radius.

GROW-CNM addresses this issue without requiring access to the original images (Adams and Bischof, 1994; Falcao et al., 2004). Lesion network mapping preserves spatial geometry through patient-specific masks,(Boes et al., 2015; Fox, 2018) but most of the published neuroimaging literature reports only cluster information (such as peak coordinates and cluster size). GROW reconstructs each cluster’s likely extent and shape by growing from the reported peak along a global, weight-ranked map of voxel-wise effects. This keeps the spatial extent and relative weighting of each finding. Our findings matched this theoretical advantage on several fronts compared with fixed-CNM: (1) Correspondence with ANM: GROW-CNM showed consistently higher correspondence with ANM, including when reference weight maps came from independent cohorts (TMUGH vs ADNI) or merge of them; (2) Robustness to study-level thresholds: GROW-CNM was less affected by the threshold used in each study (t=3.1,5,7); (3) Sampling stability: GROW-CNM was more stable under bootstrap resampling, showing higher peak probability median and lower CVs in core network.

A second issue is how heterogeneity across studies affects CNM. Cross-study confounders in literature could come from two sources: non-biological systematic bias such as scanner hardware, GMV quantification and statistical modeling, and biological sample demographic bias such as age, sex, education and disease severity (Abdulkadir et al., 2011; Fortin et al., 2018; Sauty et al., 2023). Our study found that these confounders do not act as uniform background noise but exert diverse influences on fixed-CNM and GROW-CNM, respectively. Systematic factors (statistical model, GMV type, field strength, and MRI vendor) mainly affected Fixed-CNM, including the posterior parietal, medial prefrontal, thalamic, and temporal pole regions, and could explained about 44.4% mismatch between adjusted and unadjusted fixed-CNM 50% probability map. This indicates that much of its apparent convergence reflected methodological differences between studies rather than disease effects. In contrast, GROW-CNM was associated with only one systematic factor (statistical model) and retained 93.1% (6.9% mismatch) overlap after systematic adjustment. Because systematic factors carry no biological meaning, they should routinely be adjusted for in CNM, and GROW-CNM was substantially less affected by them.

Demographic factors showed the opposite pattern. Education, MMSE score, and male ratio were associated with GROW-CNM but not with Fixed-CNM. One possible explanation is that fixed-CNM may be insensitive to demographic factors rather than robust to them. Because these associations may reflect biological factors such as disease severity, cognitive reserve, or sex, demographic adjustment involves a trade-off. It reduces bias from differences in sampling, but it may also remove meaningful disease-related variation. Demographic factors may therefore be better examined as moderators than simply removed. Because they are study-level averages, they also cannot be interpreted as effects in individual patients.

Besides, this confound burden was method-dependent: for GROW-CNM, high-probability core nodes were markedly more resilient to confounders than peripheral region, whereas for fixed-radius CNM the reverse held, with confounds concentrated within the core network. Overall, GROW-CNM is more robust to systematic confounding and preserves sample-related variation that Fixed-CNM may miss. Thus, by replacing arbitrary spheres with statistical cluster topographies, GROW-CNM both protects the core and reduces confound-driven peripheral inflations, clarifying the boundary between disease circuitry and instrumentation artifacts.

Beyond its methodological implications, this work helps clarify the network-level organization of atrophy in AD. VBM studies have reported inconsistent atrophy peaks across the medial temporal lobe, PCC, precuneus, angular gyrus, and lateral prefrontal cortex,.(Wang et al., 2015; Yang et al., 2012) and ANM was proposed to reconcile these findings within a shared default mode network (DMN) (Greicius et al., 2004; Lee et al., 2026; Seeley et al., 2009; Tetreault et al., 2020). Our results suggest that the degree of convergence depends strongly on seed modeling. Fixed-CNM produced diffuse, low-probability spread, whereas GROW-CNM integrated the same coordinates into a highly convergent DMN core (peak probability, median [IQR]: 0.99 [0.01]), indicating that the apparent inconsistency of the AD literature may partly reflect seed-modeling artefacts. This distinction matters because peripheral extensions of AD network maps are often used to explain clinical heterogeneity (e.g., atypical variants)(Lehmann et al., 2013; Warren et al., 2012) or to propose neuromodulation targets (Lin et al., 2019; Lozano et al., 2016; Siddiqi et al., 2020). In GROW-CNM, confound-related variance was concentrated in peripheral regions. In Fixed-CNM, it was concentrated within the putative core, suggesting that the conventional core may partly reflect methodological heterogeneity. Candidate targets should therefore not be selected from unadjusted Fixed-CNM maps. Besides, Peripheral GROW-CNM regions should be interpreted with caution and examined with moderator analyses, because their demographic associations may carry biological meaning. Consequently, our results caution against electing targets from unadjusted fixed-radius maps—whether core or peripheral—and against over-interpreting peripheral extensions of GROW-CNM maps without confound adjustment. GROW-CNM thus provides a confound-resilient template for modeling transneuronal tau/amyloid spreading(Braak and Braak, 1991; Franzmeier et al., 2020; Raj et al., 2012; Sepulcre et al., 2013; Vogel et al., 2023; Zhou et al., 2012) and may provide a more reliable foundation for nominating high-probability targets (e.g., the PCC/precuneus core) in future neuromodulation trials(Lin et al., 2019; Lozano et al., 2016).

These findings also apply more broadly to meta-analytic network mapping in disorders across neurological and psychiatric disorders, such as depression(Zhukovsky et al., 2021) and schizophrenia(Wang et al., 2024), where CNM is increasingly used to synthesize heterogeneous, underpowered findings. Our results indicate that fixed-radius CNM should not be adopted by default. When cluster extents or peak statistics are available, cluster-reconstruction approaches such as GROW should be preferred. When only coordinates are reported, spherical seeding should be acknowledged as a strong modeling assumption. Radius sensitivity analyses alone are insufficient, because Fixed-CNM maps were nearly identical across radii yet still diverged from ANM. Where possible, coordinate-based networks should be benchmarked against image-derived references. Finally, population-level synthesis should move from simple binarized overlays to multivariable models explicitly adjusting for systematic confounders and treating demographic factors as potential moderators(Bürkle et al., 2025; Fortin et al., 2018; Sauty et al., 2023).

Several limitations warrant consideration. First, validation was conducted in AD atrophy with a particular normative connectome; Future studies should test GROW-CNM in other diseases, imaging modalities, and connectomes(Taylor et al., 2021). Second, ANM is a reference rather than a ground-truth disease circuit; independent validation using molecular imaging or clinical outcomes is needed. Third, GROW-CNM requires accessible requires cluster information and a reference weight map, limiting its use in coordinate-only studies; more complete reporting and data sharing could improve applicability. Finally, analyses of study-level confounders were ecological and based on incomplete metadata. They therefore cannot be generalized to individual patients, and residual confounding cannot be excluded.

In conclusion, while coordinate network mapping offers a practical way to synthesize fragmented neuroimaging findings into circuit-level models(Siddiqi et al., 2022; Taylor et al., 2021), but its validity depends substantially on how seeds are represented and how studies are aggregated. Traditional fixed-radius CNM corresponded poorly with image-derived networks in AD, and changing the radius did not correct this. By reconstructing cluster morphology from reported cluster information and a reference statistical weight map, GROW-CNM improved study-level correspondence with ANM, cross-study convergence, resampling stability, and resilience to systematic confounding. Combining GROW-based seed reconstruction with confounder-adjusted population modeling may provide a more robust framework for mapping disease-related brain networks and prioritizing candidate neuromodulation targets, pending validation in other disorders and against independent benchmarks.

## Data availability

The source code for the Global weight-Ranked cluster-grOWing (GROW) algorithm and the CNM pipeline has been made publicly available on GitHub (https://github.com/ly233-hhh/GROW-CNM). The ADNI dataset analyzed during the current study is available in the ADNI repository (http://adni.loni.usc.edu). The extracted study-level coordinates and metadata from the literature search are provided in the Supplementary Material.

## Ethics statement

This study was conducted in accordance with the Declaration of Helsinki and was approved by the Ethics Committee of Tianjin Medical University General Hospital (Approval No. IRB2020-YX-161-01). Written informed consent was obtained from all participants or their legal representatives prior to participation, and their privacy rights have been strictly observed. For the Alzheimer’s Disease Neuroimaging Initiative (ADNI) cohort, data were obtained from the ADNI database in compliance with their data use agreement, and all participants provided written informed consent at the time of original data collection.

## Supporting information

PRISMA_2020_checklist

## Acknowledgments

We would like to thank the staff of the Tianjin Key Laboratory of Functional Imaging and Tianjin Medical University General Hospital for technical and administrative support. We also thank the Chinese Imaging Genetics (CHIMGEN) cohort for providing the normative connectome data, and the Alzheimer’s Disease Neuroimaging Initiative (ADNI) for providing the public datasets.

## Reference

Abdulkadir, A., Mortamet, B., Vemuri, P., Jack, C.J.r., Krueger, G., Klöppel, S., Initi, A.s.D.N., 2011. Effects of hardware heterogeneity on the performance of SVM Alzheimer’s disease classifier. Neuroimage 58, 785–792.

Adams, R., Bischof, L., 1994. Seeded region growing. IEEE Transactions on Pattern Analysis and Machine Intelligence 16, 641–647.

Boes, A.D., Prasad, S., Liu, H., Liu, Q., Pascual-Leone, A., Caviness, V.S., Jr., Fox, M.D., 2015. Network localization of neurological symptoms from focal brain lesions. Brain 138, 3061–3075.

Braak, H., Braak, E., 1991. Neuropathological stageing of Alzheimer-related changes. Acta neuropathologica 82, 239–259.

Bürkle, E., Nazzal, A., Debolski, A., Ernemann, U., Lindig, T., Bender, B., 2025. Scan-rescan reliability assessment of brain volumetric analysis across scanners and software solutions. Scientific Reports 15.

Darby, R.R., Joutsa, J., Fox, M.D., 2019. Network localization of heterogeneous neuroimaging findings. Brain 142, 70–79.

Dong, H.Y., Guo, L.N., Yang, H.L., Zhu, W.S., Liu, F., Xie, Y.Y., Zhang, Y., Xue, K.Z., Li, Q., Liang, M., Zhang, N., Qin, W., 2023. Association between gray matter atrophy, cerebral hypoperfusion, and cognitive impairment in Alzheimer’s disease. Frontiers in Aging Neuroscience 15.

Dubois, B., Feldman, H.H., Jacova, C., 2014. Advancing research diagnostic criteria for Alzheimer’s disease: the IWG-2 criteria (vol 13, pg 614, 2014). Lancet Neurology 13, 757–757.

Eickhoff, S.B., Bzdok, D., Laird, A.R., Kurth, F., Fox, P.T., 2012. Activation likelihood estimation meta-analysis revisited. Neuroimage 59, 2349–2361.

Falcao, A.X., Stolfi, J., Lotufo, R.D.A., 2004. The image foresting transform: Theory, algorithms, and applications. Ieee Transactions on Pattern Analysis and Machine Intelligence 26, 19–29.

Fortin, J.P., Cullen, N., Sheline, Y.I., Taylor, W.D., Aselcioglu, I., Cook, P.A., Adams, P., Cooper, C., Fava, M., McGrath, P.J., McInnis, M., Phillips, M.L., Trivedi, M.H., Weissman, M.M., Shinohara, R.T., 2018. Harmonization of cortical thickness measurements across scanners and sites. Neuroimage 167, 104–120.

Fox, M.D., 2018. Mapping Symptoms to Brain Networks with the Human Connectome. N Engl J Med 379, 2237–2245.

Franzmeier, N., Neitzel, J., Rubinski, A., Smith, R., Strandberg, O., Ossenkoppele, R., Hansson, O., Ewers, M., Adni, 2020. Functional brain architecture is associated with the rate of tau accumulation in Alzheimer’s disease. Nature Communications 11.

Greicius, M.D., Srivastava, G., Reiss, A.L., Menon, V., 2004. Default-mode network activity distinguishes Alzheimer’s disease from healthy aging: Evidence from functional MRI. Proceedings of the National Academy of Sciences of the United States of America 101, 4637–4642.

Ji, G.J., Xu, W.Q., Zhang, J., Li, Q.Q., Yang, Y.N., He, K.L., Yu, F.Q., Zhu, C.Y., Jiang, X., Jing, L., Hu, P.P., Tian, Y.H., Wang, K., 2026. Distinct antidepressant therapies act on a common brain network. Nature Communications 17.

Lam, B., Masellis, M., Freedman, M., Stuss, D.T., Black, S.E., 2013. Clinical, imaging, and pathological heterogeneity of the Alzheimer’s disease syndrome. Alzheimers Research & Therapy 5.

Lee, S., Baratono, S.R., Ha, J., Burt, G.T., Palm, S.T., Drew, W.J., Zide, B.S., Chiulli, N.M., Lariviere, S., Zhang, S.S., Yeo, B.T.T., Fox, M.D., Sperling, R.A., Donovan, N.J., Siddiqi, S.H., 2026. Atrophy in preclinical Alzheimer’s disease maps to a network that predicts longitudinal decline. Molecular Psychiatry.

Lehmann, M., Madison, C.M., Ghosh, P.M., Seeley, W.W., Mormino, E., Greicius, M.D., Gorno-Tempini, M.L., Kramer, J.H., Miller, B.L., Jagust, W.J., Rabinovici, G.D., 2013. Intrinsic connectivity networks in healthy subjects explain clinical variability in Alzheimer’s disease. Proceedings of the National Academy of Sciences of the United States of America 110, 11606–11611.

Lin, Y., Jiang, W.J., Shan, P.Y., Lu, M., Wang, T., Li, R.H., Zhang, N., Ma, L., 2019. The role of repetitive transcranial magnetic stimulation (rTMS) in the treatment of cognitive impairment in patients with Alzheimer’s disease: A systematic review and meta-analysis. Journal of the Neurological Sciences 398, 184–191.

Lozano, A.M., Fosdick, L., Chakravarty, M.M., Leoutsakos, J.M., Munro, C., Oh, E., Drake, K.E., Lyman, C.H., Rosenberg, P.B., Anderson, W.S., Tang-Wai, D.F., Pendergrass, J.C., Salloway, S., Asaad, W.F., Ponce, F.A., Burke, A., Sabbagh, M., Wolk, D.A., Baltuch, G., Okun, M.S., Foote, K.D., McAndrews, M.P., Giacobbe, P., Targum, S.D., Lyketsos, C.G., Smith, G.S., 2016. A Phase II Study of Fornix Deep Brain Stimulation in Mild Alzheimer’s Disease. Journal of Alzheimers Disease 54, 777–787.

Menon, V., 2011. Large-scale brain networks and psychopathology: a unifying triple network model. Trends Cogn Sci 15, 483–506.

Page, M.J., McKenzie, J.E., Bossuyt, P.M., Boutron, I., Hoffmann, T.C., Mulrow, C.D., Shamseer, L., Tetzlaff, J.M., Akl, E.A., Brennan, S.E., Chou, R., Glanville, J., Grimshaw, J.M., Hrobjartsson, A., Lalu, M.M., Li, T.J., Loder, E.W., Mayo-Wilson, E., McDonald, S., McGuinness, L.A., Stewart, L.A., Thomas, J., Tricco, A.C., Welch, V.A., Whiting, P., Moher, D., 2021. The PRISMA 2020 statement: an updated guideline for reporting systematic reviews. Bmj-British Medical Journal 372.

Paquola, C., De Wael, R.V., Wagstyl, K., Bethlehem, R.A.I., Hong, S.J., Seidlitz, J., Bullmore, E.T., Evans, A.C., Misic, B., Margulies, D.S., Smallwood, J., Bernhardt, B.C., 2019. Microstructural and functional gradients are increasingly dissociated in transmodal cortices. Plos Biology 17.

Petersen, R.C., Aisen, P.S., Beckett, L.A., Donohue, M.C., Gamst, A.C., Harvey, D.J., Jack, C.R., Jagust, W.J., Shaw, L.M., Toga, A.W., Trojanowski, J.Q., Weiner, M.W., 2010. Alzheimer’s Disease Neuroimaging Initiative (ADNI) Clinical characterization. Neurology 74, 201–209.

Raj, A., Kuceyeski, A., Weiner, M., 2012. A Network Diffusion Model of Disease Progression in Dementia. Neuron 73, 1204–1215.

Sauty, B., Durrleman, S., Alzheimers Dis, N., 2023. Impact of sex and APOE-ε4 genotype on patterns of regional brain atrophy in Alzheimer’s disease and healthy aging. Frontiers in Neurology 14.

Schaefer, A., Kong, R., Gordon, E.M., Laumann, T.O., Zuo, X.N., Holmes, A.J., Eickhoff, S.B., Yeo, B.T.T., 2018. Local-Global Parcellation of the Human Cerebral Cortex from Intrinsic Functional Connectivity MRI. Cerebral Cortex 28, 3095–3114.

Seeley, W.W., Crawford, R.K., Zhou, J., Miller, B.L., Greicius, M.D., 2009. Neurodegenerative diseases target large-scale human brain networks. Neuron 62, 42–52.

Sepulcre, J., Sabuncu, M.R., Becker, A., Sperling, R., Johnson, K.A., 2013. characterization of the early states of the amyloid-beta network. Brain 136, 2239–2252.

Shen, Y.H., Cai, H.H., Mo, F., Yao, S.W., Yu, Y.Q., Zhu, J.J., 2023. Functional connectivity gradients of the cingulate cortex. Communications Biology 6.

Siddiqi, S.H., Kording, K.P., Parvizi, J., Fox, M.D., 2022. Causal mapping of human brain function. Nature Reviews Neuroscience 23, 361–375.

Siddiqi, S.H., Taylor, S.F., Cooke, D., Pascual-Leone, A., George, M.S., Fox, M.D., 2020. Distinct Symptom-Specific Treatment Targets for Circuit-Based Neuromodulation. American Journal of Psychiatry 177, 435–446.

Taylor, J.J., Siddiqi, S.H., Fox, M.D., 2021. Coordinate Network Mapping: An Emerging Approach for Morphometric Meta-Analysis. American Journal of Psychiatry 178, 1080–1081.

Tetreault, A.M., Phan, T., Orlando, D., Lyu, I., Kang, H., Landman, B., Darby, R.R., Initia, A.D.N., 2020. Network localization of clinical, cognitive, and neuropsychiatric symptoms in Alzheimer’s disease. Brain 143, 1249–1260.

Vogel, J.W., Corriveau-Lecavalier, N., Franzmeier, N., Pereira, J.B., Brown, J.A., Maass, A., Botha, H., Seeley, W.W., Bassett, D.S., Jones, D.T., Ewers, M., 2023. Connectome-based modelling of neurodegenerative diseases: towards precision medicine and mechanistic insight. Nature Reviews Neuroscience 24, 620–639.

Wang, W.Y., Yu, J.T., Liu, Y., Yin, R.H., Wang, H.F., Wang, J., Tan, L., Radua, J., Tan, L., 2015. Voxel-based meta-analysis of grey matter changes in Alzheimer’s disease. Translational Neurodegeneration 4.

Wang, Y.R., Yang, Y.N., Xu, W.Q., Yao, X.Q., Xie, X.H., Zhang, L., Sun, J.M., Wang, L., Hua, Q., He, K.L., Tian, Y.H., Wang, K., Ji, G.J., 2024. Heterogeneous Brain Abnormalities in Schizophrenia Converge on a Common Network Associated With Symptom Remission. Schizophrenia Bulletin 50, 545–556.

Warren, J.D., Fletcher, P.D., Golden, H.L., 2012. The paradox of syndromic diversity in Alzheimer disease. Nature Reviews Neurology 8, 451–464.

Xu, Q., Guo, L.N., Cheng, J.L., Wang, M.Y., Geng, Z.J., Zhu, W.Z., Zhang, B., Liao, W.H., Qiu, S.J., Zhang, H., Xu, X.J., Yu, Y.Q., Gao, B., Han, T., Yao, Z.W., Cui, G.B., Liu, F., Qin, W., Zhang, Q., Li, M.L.J., Liang, M., Chen, F., Xian, J.F., Li, J.C., Zhang, J., Zuo, X.N., Wang, D.W., Shen, W., Miao, Y.W., Yuan, F., Lui, S., Zhang, X.C., Xu, K., Zhang, L.J., Ye, Z.X., Yu, C.S., Consortium, C., 2020. CHIMGEN: a Chinese imaging genetics cohort to enhance cross-ethnic and cross-geographic brain research. Molecular Psychiatry 25, 517–529.

Yang, J., Pan, P.L., Song, W., Huang, R., Li, J.P., Chen, K., Gong, Q.Y., Zhong, J.G., Shi, H.C., Shang, H.F., 2012. Voxelwise meta-analysis of gray matter anomalies in Alzheimer’s disease and mild cognitive impairment using anatomic likelihood estimation. Journal of the Neurological Sciences 316, 21–29.

Yarkoni, T., Poldrack, R.A., Nichols, T.E., Van Essen, D.C., Wager, T.D., 2011. Large-scale automated synthesis of human functional neuroimaging data. Nature Methods 8, 665–U695.

Yeo, B.T.T., Krienen, F.M., Sepulcre, J., Sabuncu, M.R., Lashkari, D., Hollinshead, M., Roffman, J.L., Smoller, J.W., Zöller, L., Polimeni, J.R., Fischl, B., Liu, H.S., Buckner, R.L., 2011. The organization of the human cerebral cortex estimated by intrinsic functional connectivity. Journal of Neurophysiology 106, 1125–1165.

Zhou, J., Gennatas, E.D., Kramer, J.H., Miller, B.L., Seeley, W.W., 2012. Predicting Regional Neurodegeneration from the Healthy Brain Functional Connectome. Neuron 73, 1216–1227.

Zhukovsky, P., Anderson, J.A.E., Coughlan, G., Mulsant, B.H., Cipriani, A., Voineskos, A.N., 2021. Coordinate-Based Network Mapping of Brain Structure in Major Depressive Disorder in Younger and Older Adults: A Systematic Review and Meta-Analysis. American Journal of Psychiatry 178, 1119–1128.

