## Supplementary figures for "Global weight-Ranked cluster-grOWing (GROW): A Cluster-Growing Framework for Robust Functional Coordinate Network Mapping"

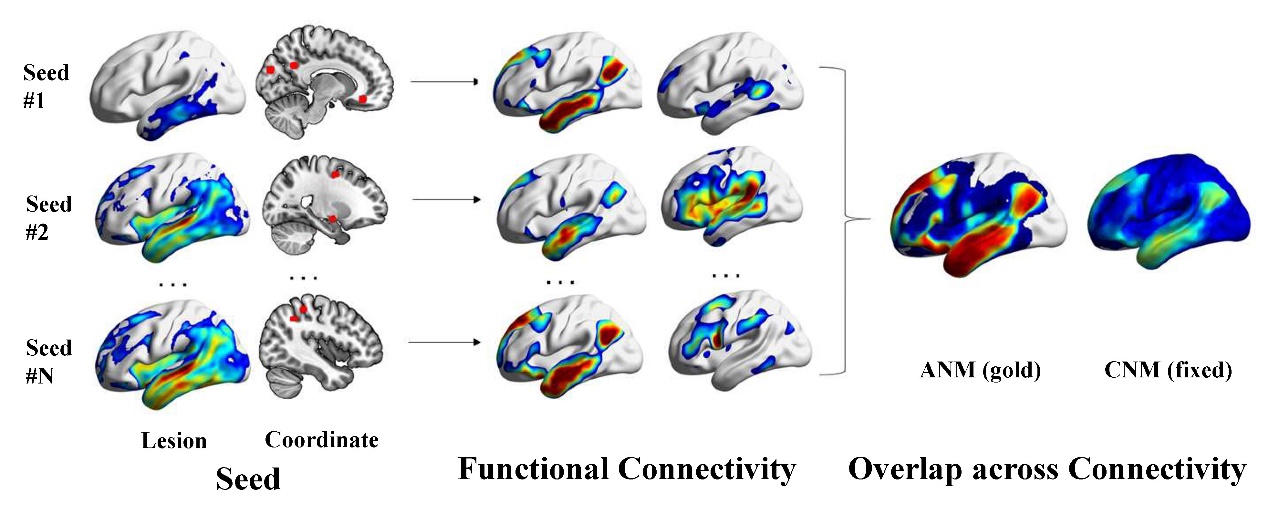

**Supplementary Figure 1. Study-level functional network mapping pipeline.** Schematic illustration of the study-level ANM procedure. For each seed region (atrophy cluster for ANM and fixed-radius sphere for fixed-CNM,), the mean resting-state time series of each seed were extracted from the normative connectome and correlated with all other voxels to generate a seed-to-voxel functional connectivity map. Individual connectivity maps were then transformed to Fisher's z and averaged across participants to produce population-level spatial overlap maps. ANM, Atrophy Network Mapping; CNM, Coordinate Network Mapping.

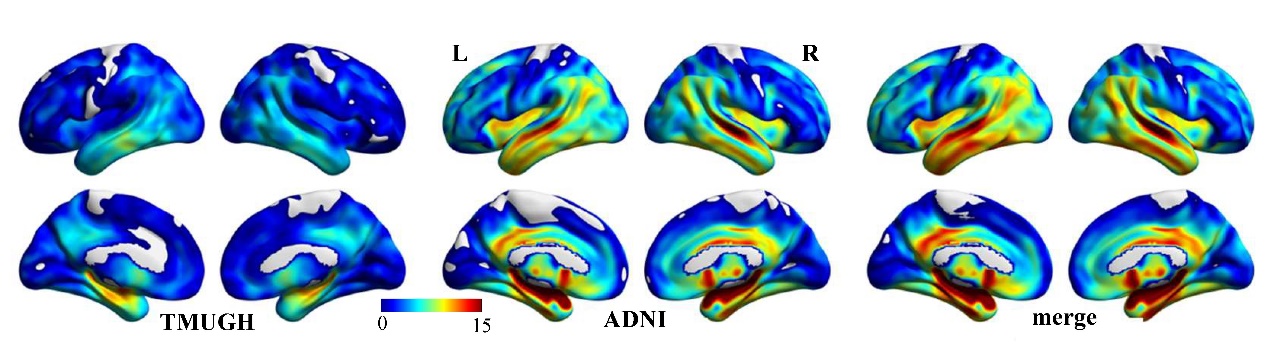

**Supplementary Figure 2**. **Unthresholded t-statistic maps of GMV atrophy of Alzheimer's disease patients.** The color bar represents t-values (warm colors indicate AD < NC). AD, Alzheimer’s disease; NC, Normal Controls.

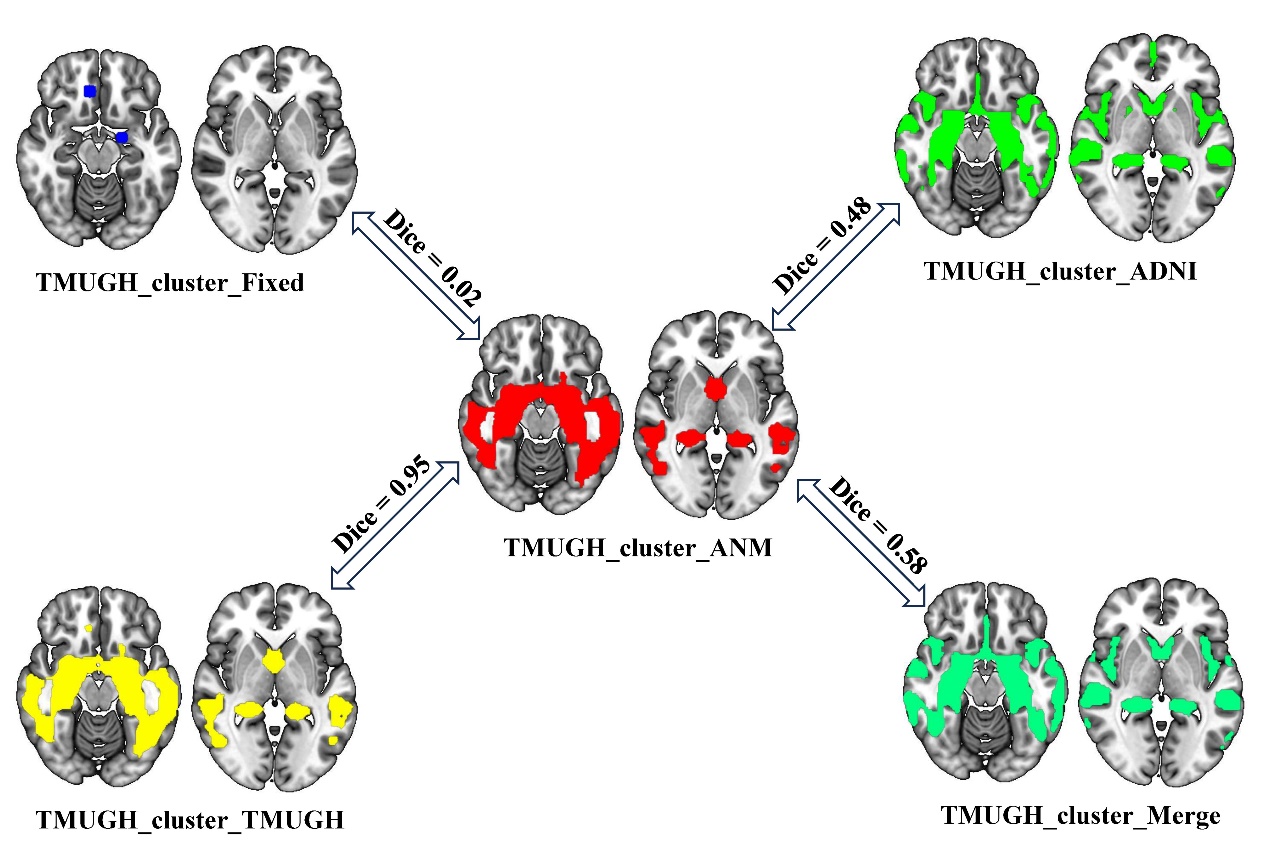

**Supplementary Figure 3. Spatial overlap between observed atrophy clusters and GROW-reconstructed clusters in the TMUGH cohort using different reference weights.** The panels show binary fixed-radius seeds (upper left) and GROW-reconstructed clusters using t-maps from the TMUGH (upper right), ADNI (lower left), and merged datasets (lower right) as reference weight maps. The center panel shows the observed atrophy clusters in the TMUGH dataset (P < 0.05, family-wise error corrected). Numbers indicate Dice coefficients relative to the observed clusters. ANM, Atrophy Network Mapping; ADNI, Alzheimer’s Disease Neuroimaging Initiative; TMUGH, Tianjin Medical University General Hospital.

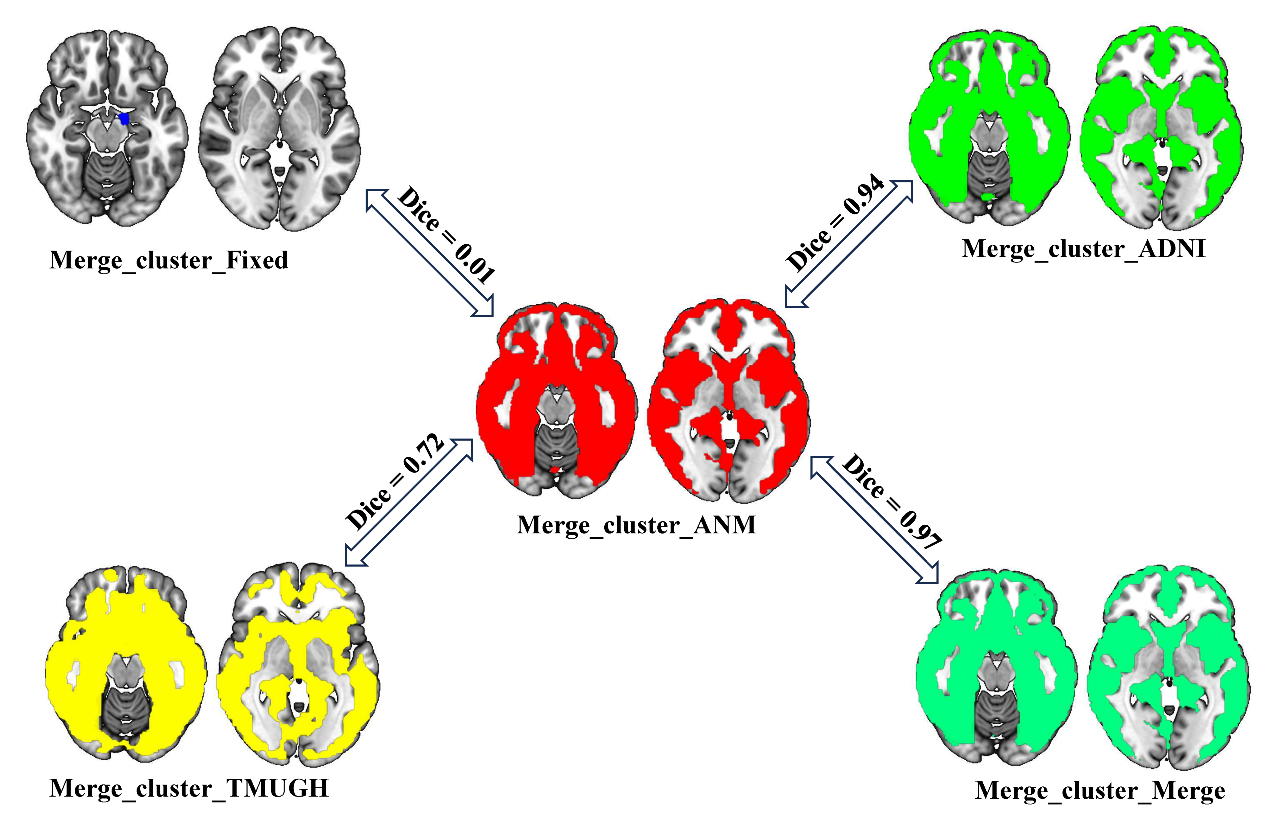

**Supplementary Figure 4. Spatial overlap between observed atrophy clusters and GROW-reconstructed clusters in the merge cohort using different reference weights.** The panels show binary fixed-radius seeds (upper left) and GROW-reconstructed clusters using t-maps from the TMUGH (upper right), ADNI (lower left), and merged datasets (lower right) as reference weight maps. The center panel shows the observed atrophy clusters in the TMUGH dataset (P < 0.05, family-wise error corrected). Numbers indicate Dice coefficients relative to the observed clusters. ANM, Atrophy Network Mapping; ADNI, Alzheimer’s Disease Neuroimaging Initiative; TMUGH, Tianjin Medical University General Hospital.

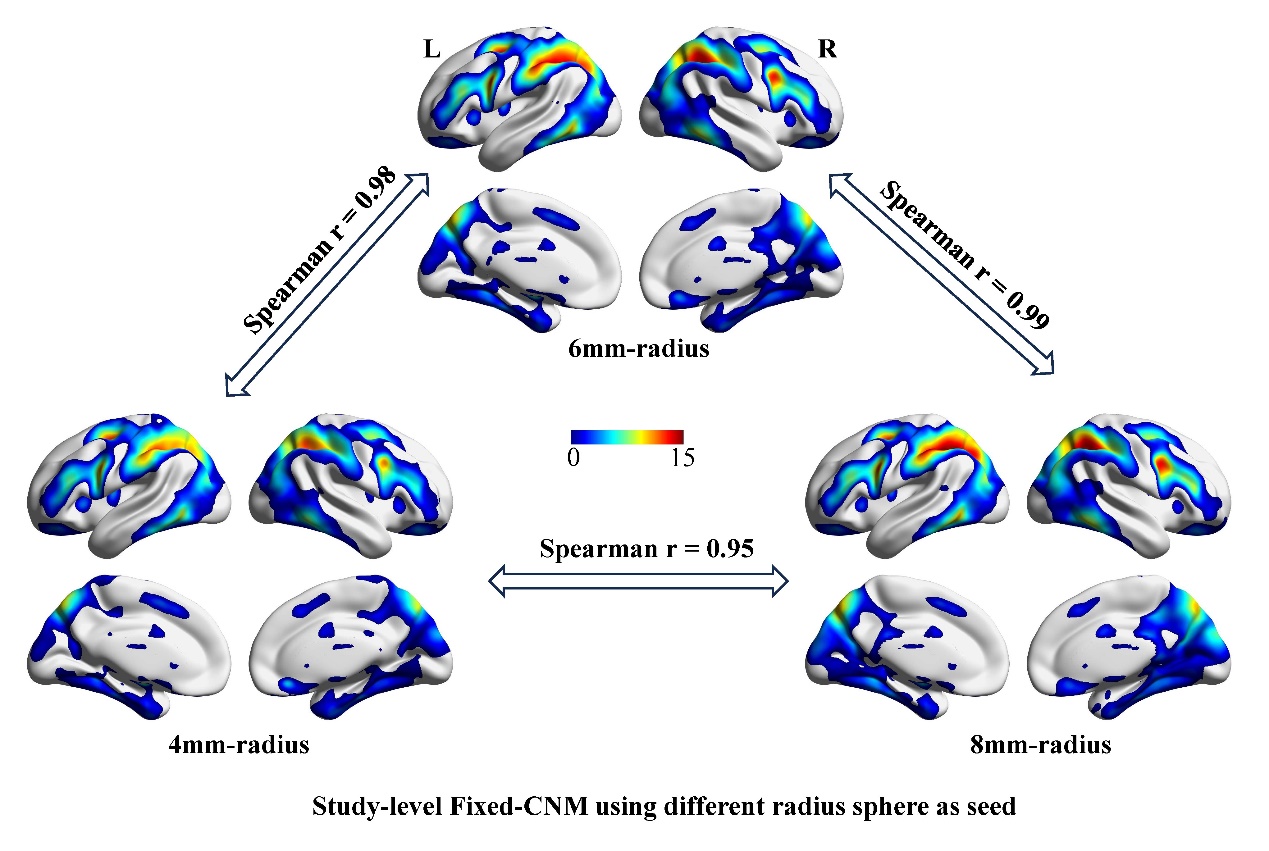

**Supplementary Figure 5.** **Sensitivity analysis of seed radius in Fixed-CNM.** Study-level functional connectivity maps were generated using spherical seeds centered at the reported peak coordinates with three different radii: 4 mm, 6 mm, and 8 mm. The color bar represents t value of functional connectivity between the seed region and each voxel in the whole brain. The overall spatial patterns—particularly the involvement of the default mode network regions—are highly consistent across the three radius conditions, confirming that the Fixed-CNM network topography is not critically dependent on the specific choice of seed radius within this range. CNM, Coordinate Network Mapping.

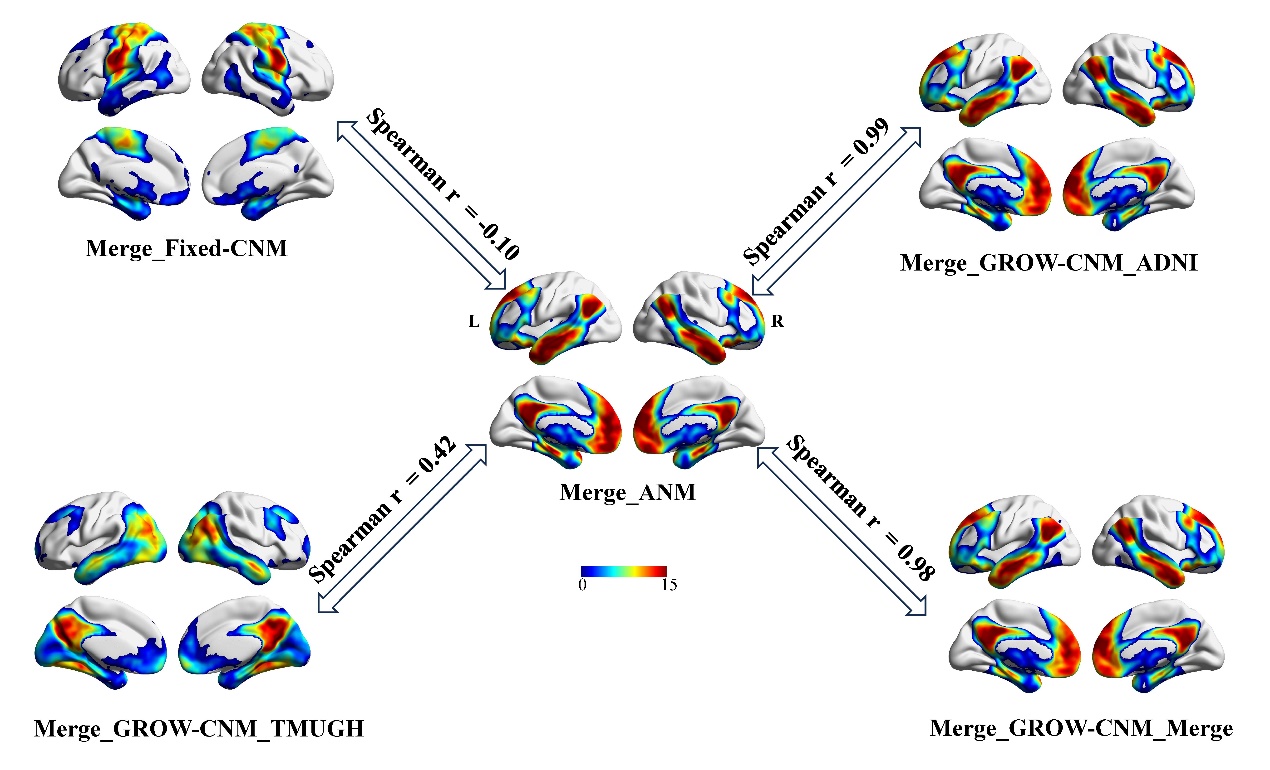

**Supplementary Figure 6. Study-level functional connectivity maps from different CNM approaches compared with the reference ANM-derived map in the Merge cohort.** Functional connectivity maps were generated using seeds derived from Fixed-CNM (upper left) and GROW algorithm with different weight maps—ADNI cluster weight map (upper right), TMUGH cluster weight map (lower left), and merged weight map (lower right)—alongside the reference ANM-derived map (center). Spearman’s r values indicate the spatial correlation between each CNM-derived map and the reference ANM map. CNM, Coordinate Network Mapping; ANM, Atrophy Network Mapping; ADNI, Alzheimer's Disease Neuroimaging Initiative; TMUGH, Tianjin Medical University General Hospital; GROW, Global weight-Ranked cluster-grOWing.

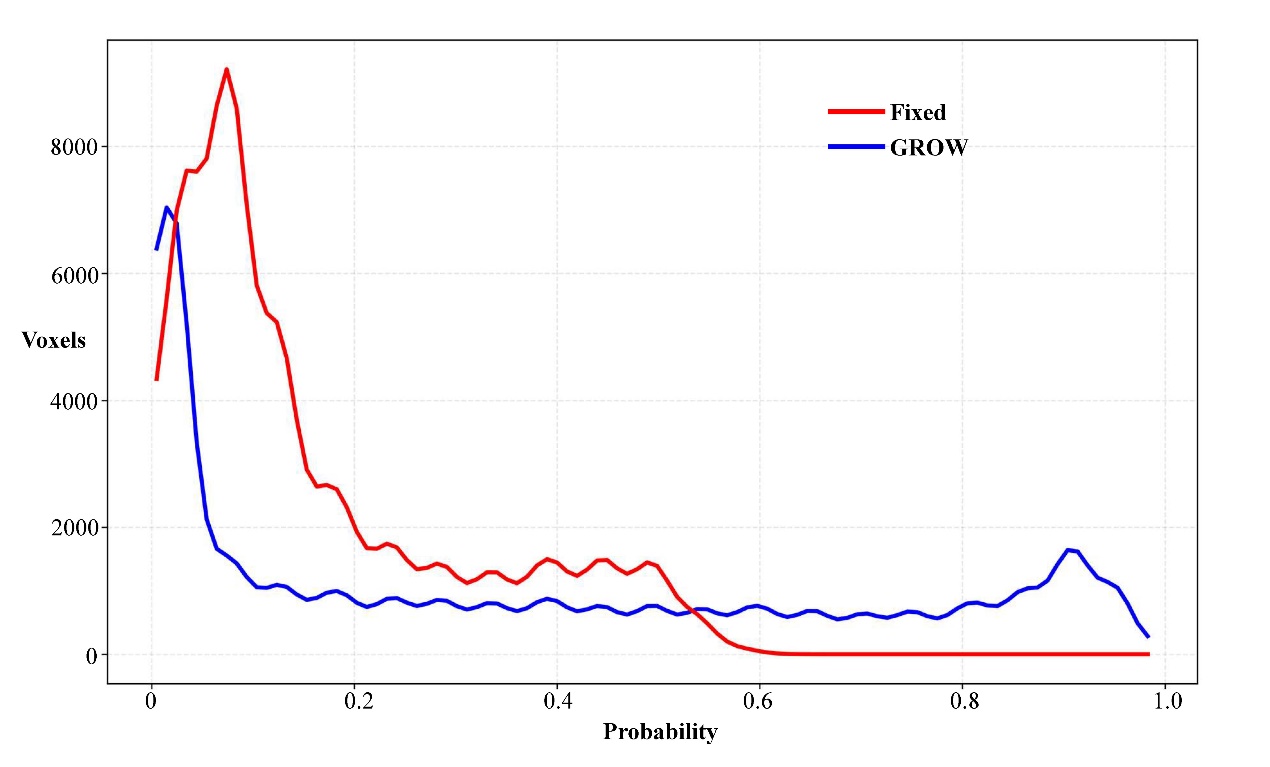

**Supplementary Figure 7. Histogram of probability map for Fixed-CNM and GROW-CNM.** The x-axis shows probability thresholds from 0 to 1, and the y-axis shows the number of voxels (in thousands). Red and blue lines represent Fixed-CNM and GROW-CNM, respectively. Maps were derived from unadjusted population-level probability maps. CNM, Coordinate Network Mapping; GROW, Global weight-Ranked cluster-grOWing.

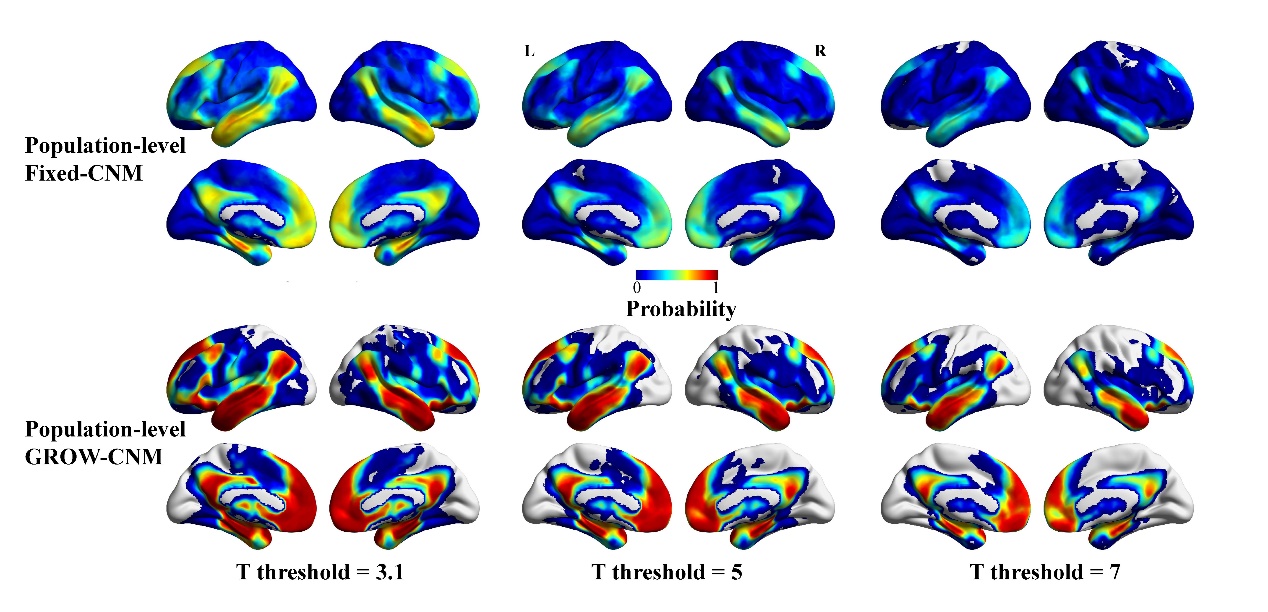

**Supplementary Figure 8. Population-level Fixed-CNM and GROW-CNM maps across study-level thresholds.** Study-level functional connectivity maps were thresholded at t = 3.1, 5, or 7, binarized, and combined to generate population-level probability maps. The color bar indicates the proportion of studies with suprathreshold functional connectivity at each voxel. CNM, Coordinate Network Mapping; GROW, Global weight-Ranked cluster-grOWing.

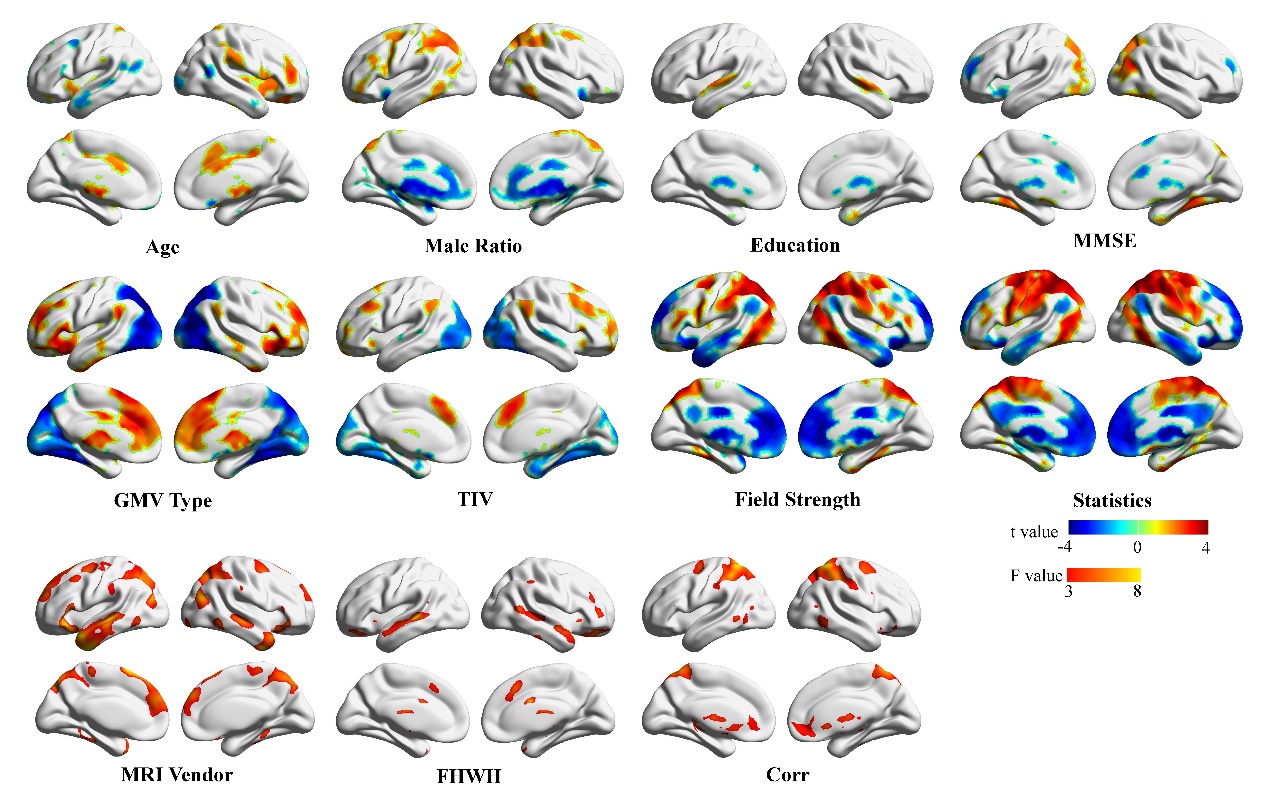

**Supplementary Figure 9. Effect of single confounder on fixed-CNM.** Voxel-wise general linear models were performed to evaluate the association between single systematic or demographic factor on the study-level fixed-CNM maps (p < 0.05, uncorrected). T-statistic and F-statistic maps are presented separately according to the factor type. MMSE, Mini-Mental State Examination; GMV, Gray Matter Volume; TIV, Total Intracranial Volume; MRI, Magnetic Resonance Imaging; FWHM, Full-Width at Half-Maximum; Corr, Correction.

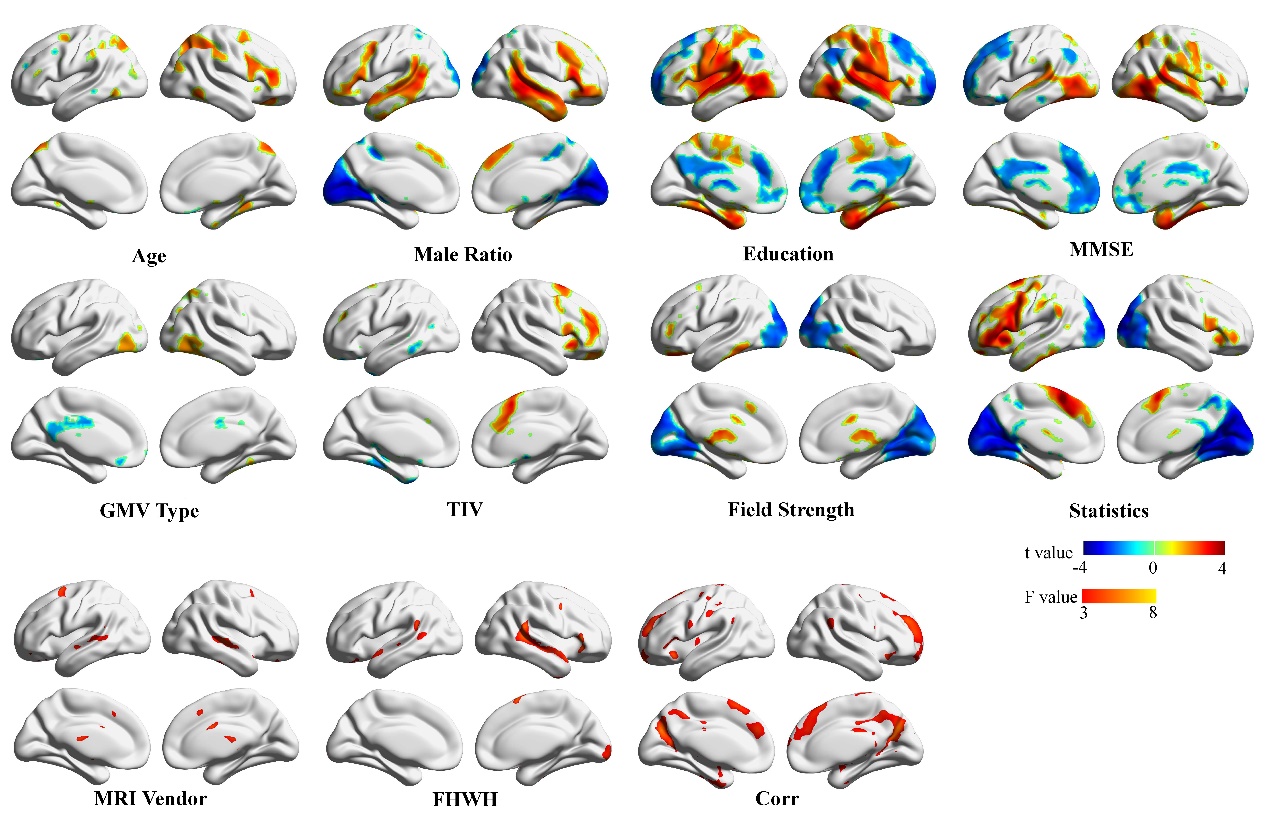

**Supplementary Figure 10. Effect of single confounder on GROW-CNM.** Voxel-wise general linear models were performed to evaluate the association between single systematic or demographic factor on the study-level GROW-CNM maps (p < 0.05, uncorrected). T-statistic and F-statistic maps are presented separately according to the factor type. MMSE, Mini-Mental State Examination; GMV, Gray Matter Volume; TIV, Total Intracranial Volume; MRI, Magnetic Resonance Imaging; FWHM, Full-Width at Half-Maximum; Corr, Correction.

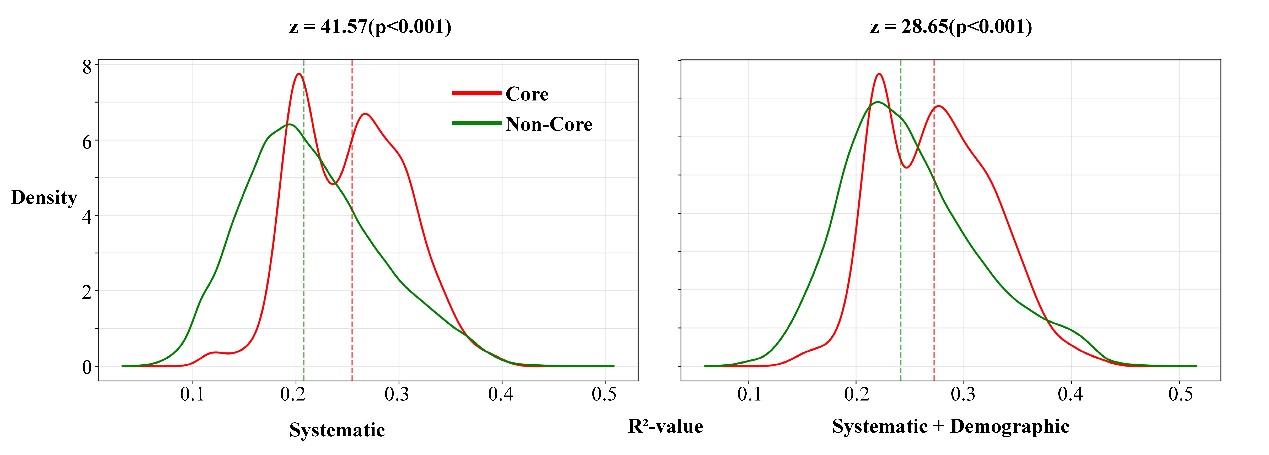

**Supplementary Figure 11. Comparison in explained confounder variance between core and non-core regions using Fixed-CNM-defined core mask.** R² values (proportion of variance explained by systematic and demographic confounders) were compared between core and non-core regions, with the core region defined as voxels with probability > 0.5 in the unadjusted Fixed-CNM probability map. For Fixed-CNM, R² values remained significantly higher in the core region than in the non-core region (p < 0.001), indicating that the concentration of confounder effects within its putative core network was not an artifact of using GROW-CNM-derived core masks. CNM, Coordinate Network Mapping.
